# A SABATH family enzyme regulates development via the gibberellin-related pathway in the liverwort *Marchantia polymorpha*

**DOI:** 10.64898/2025.12.11.693594

**Authors:** Shogo Kawamura, Eita Shimokawa, Maika Ito, Isuzu Nakamura, Takehiko Kanazawa, Megumi Iwano, Rui Sun, Yoshihiro Yoshitake, Shohei Yamaoka, Shinjiro Yamaguchi, Takashi Ueda, Misako Kato, Takayuki Kohchi

## Abstract

The SABATH family enzymes are a group of plant-specific methyltransferases that catalyze the methylation of many small molecules, including several plant hormones. While this family emerged anciently before the evolution of land plants from streptophyte algae, little is known about their biological function in plant lineages other than angiosperms. Here, we identified 12 SABATH family genes from the liverwort *Marchantia polymorpha* and found that Mp*SABATH2* is essential for its development. Mp*sabath2* mutants were severely inhibited in thallus growth and gemma cup formation, while spontaneously forming sexual branches under non-inductive conditions. These phenotypes resembled the developmental responses to far-red light, which was also supported by transcriptome analysis. Further genetic analysis connected this phenomenon with gibberellin (GA)-related metabolism. Blocking GA biosynthesis partially rescued Mp*sabath2* phenotypes, which were restored by treatment with the GA precursor, *ent*-kaurenoic acid. Given that bryophyte and angiosperm SABATH proteins fall into distinct phylogenetic clades, our findings suggest that SABATH family enzymes independently acquired roles in developmental regulation through convergent or parallel evolution in land plants.

## Introduction

Throughout evolution, plants established diverse metabolic pathways to synthesize a wide variety of specialized chemical compounds crucial for growth regulation, defense against pathogens and adaptation to abiotic stresses. Methylation, i.e., the addition of a methyl group to the substrate, is a common biochemical modification that changes the bioactivity of many plant metabolites. Such modification is mainly catalyzed by methyltransferases (MTs) that uses *S*-adenosyl-L-methionine (SAM) as a methyl donor. In plants, the SABATH family of SAM-dependent MTs is specifically associated with plant hormones and other specialized metabolites (D’Auria et al., 2003; Ward et al., 2021; Wang et al., 2024).

With the family name coined from salicylic acid carboxyl methyltransferase (SAMT), benzoic acid carboxyl methyltransferase (BAMT) and theobromine synthase, the biochemical and physiological functions of many SABATH enzymes have been characterized in flowering plants (Wang et al., 2024). Xanthosine methyltransferase, theobromine synthase, and caffeine synthase catalyze three successive methylation steps in the biosynthesis of theobromine and caffeine (Kato and Mizuno, 2004), which evolved convergently in *Coffea* (coffee), *Theobroma* (cacao), and *Camellia* (tea) species (Denoeud et al., 2014). SAMT and jasmonic acid methyltransferase (JMT) catalyze the methylation of two defense-related hormones (Ross et al., 1999; Seo et al., 2001), producing volatile products that contribute to both floral fragrance and defense responses, partly by serving as airborne signals that prime immune responses in neighboring plants (Farmer and Ryan, 1990; Shulaev et al., 1997; Ross et al., 1999; Seo et al., 2001; Koo et al., 2007). Indole-3-acetic acid carboxyl methyltransferase (IAMT) deactivates the growth hormone auxin by converting indole-3-acetic acid (IAA) to methyl-IAA, which regulates leaf morphology, gravitropism, and heat tolerance (Qin et al., 2005; Abbas et al., 2018). Similarly, gibberellin methyltransferase (GAMT) catalyzes the deactivation of gibberellins, thereby modulating seed germination and flowering time in Arabidopsis (Varbanova et al., 2007; Xing et al., 2007; Lee et al., 2020). Recently, a carlactonoic acid methyltransferase (CLAMT) was identified as a key enzyme in the biosynthesis of strigolactones, the hormone that inhibits shoot branching (Wakabayashi et al., 2021; Mashiguchi et al., 2022).

A previous phylogenetic analysis showed that SABATH enzymes broadly exist in streptophyte algae and land plants (Wang et al., 2024). However, only a few studies have examined the function of this family outside of angiosperms. In gymnosperms, enzyme activities of IAMT, SAMT, JMT, and GAMT have been reported in *Picea abies* and other species (Zhao et al., 2009; Chaiprasongsuk et al., 2018; Zhang et al., 2020). In the moss *Physcomitrium patens*, one of the four SABATH enzymes showed activity to catalyze *S*-methylation of thiols (Zhao et al., 2012). In the liverwort *Conocephalum salebrosum*, the activities of CsSAMT and cinnamic acid methyltransferase (CsCAMT) have been confirmed (Zhang et al., 2019). Meanwhile, neither IAMT, JMT nor GAMT has been found in bryophytes.

Land plants emerged from streptophyte algae around 500 million years ago and soon diverged into bryophytes and vascular plants (Morris et al., 2018; Harris et al., 2022). Such ancient evolutionary divergence has led to both conservation and diversification of plant hormone metabolic pathways between these plant lineages. While auxin, cytokinin, and abscisic acid (ABA) biosynthetic pathways are well-conserved among all land plants (Morffy and Strader, 2020; Li et al., 2022; Azar et al., 2025), jasmonic acid (JA) and gibberellin (GA) pathways are diverged between bryophytes and vascular plants. In the case of JA, jasmonoyl-isoleucine (JA-Ile) is the major bioactive JA in flowering plants (Staswick and Tiryaki, 2004), whereas the JA-Ile precursor dinor-12-oxo-phytodienoic acid (dn-OPDA) serves as the ligand to trigger JA signaling in the liverwort *Marchantia polymorpha* and other bryophytes (Monte et al., 2018; Inagaki et al., 2021; Chini et al., 2023). Similarly, bryophytes do not synthesize GAs bioactive in vascular plants (e.g. GA_1_ or GA_4_), but utilize the GA precursor, *ent*-kaurenoic acid (KA) to generate lineage-specific bioactive molecules (Hayashi et al., 2010; Miyazaki et al., 2018; Sun et al., 2023). Distinct from the growth-promoting activity of GA in angiosperms, GA-related pathway modulates far-red light (FR) responses in the liverwort *M. polymorpha*, in a manner that restricts the overall plant growth (Sun et al., 2023).

In this research, we investigated SABATH family enzymes in the liverwort *M. polymorpha.* One of these enzymes, MpSABATH2, was found to be essential for thallus growth and suppression of sexual reproduction, which is partially dependent on the biosynthesis of GA-related compounds. These findings revealed the functional importance of SABATH2 enzymes in the bryophyte lineage, providing insights into the evolution of specialized metabolism in land plants.

## Results

### SABATH family genes from the liverwort *Marchantia polymorpha*

To investigate SABATH family genes in *M. polymorpha*, we searched the standard reference genome version 7.1 (MpTak_v7.1) (Tanizawa et al., 2025) with HMMER for SAM-dependent carboxyl methyltransferases using the PFAM profile PF03492. We identified 12 SABATH family genes located on five chromosomes: five on chromosome 5, three on chromosome 4, and one each on chromosomes 2, 3, 6, and 7 (Fig. 1A; Suppl. Table S1). Three genes on chromosome 5 (Mp*SABATH7*/Mp5g24210, Mp*SABATH11*/Mp5g24220, and Mp*SABATH12*/Mp5g24250) and two genes on chromosome 4 (Mp*SABATH4*/Mp4g15440 and Mp*SABATH6*/Mp4g15450) are arranged in tandem repeats.

**Figure 1.**
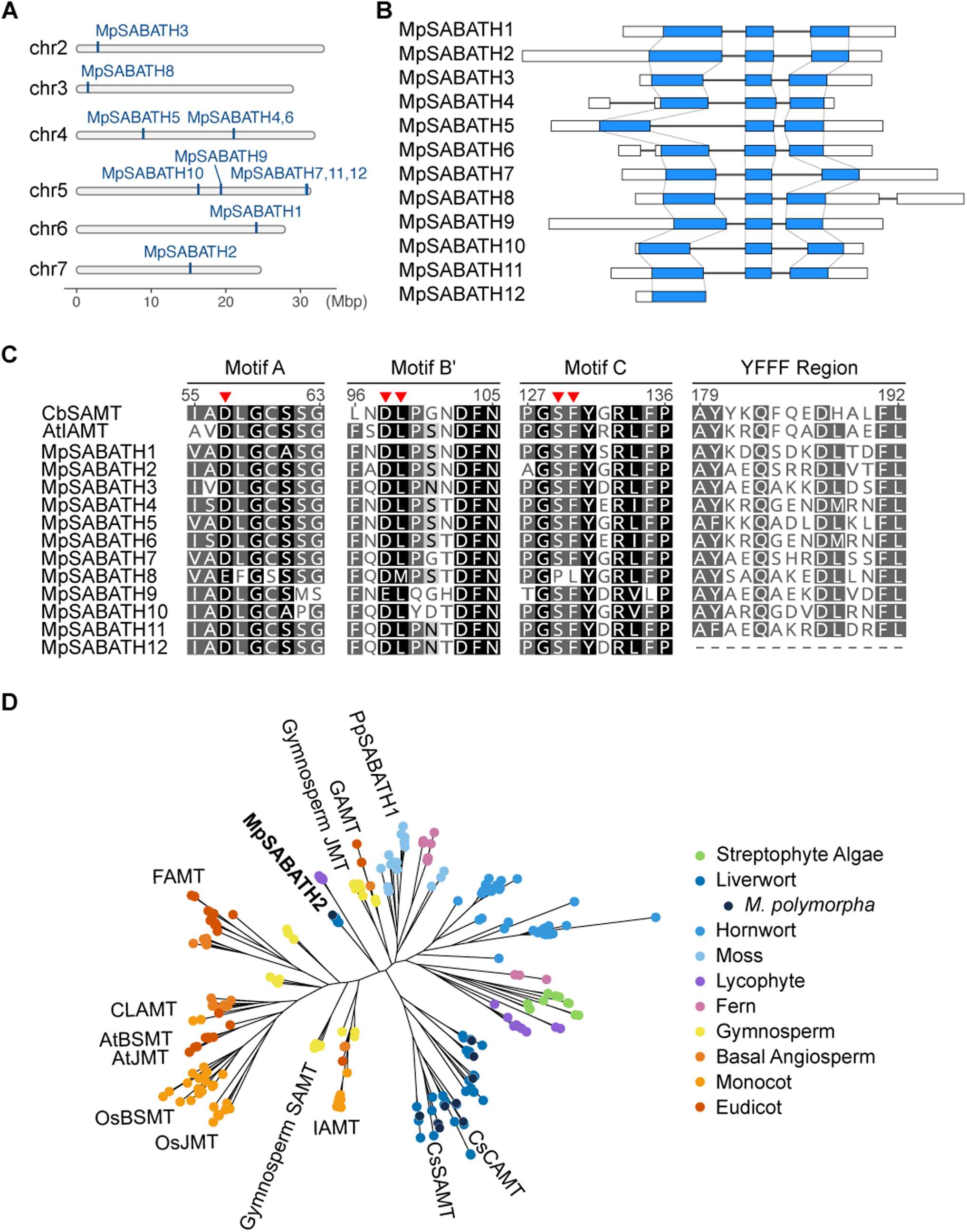
Overview of SABATH family genes in *M. polymorpha*. A, Genomic localization of Mp*SABATH* genes. Blue lines indicate the positions of Mp*SABATH* genes on each chromosome. B, Exon-intron structure of Mp*SABATH* genes. White and blue blocks represent untranslated and coding regions of exons, respectively. Black lines connecting the colored blocks represent introns. The size of the 5′ untranslated region in Mp*SABATH8* was manually curated (see also Suppl.Fig.3). C, Multiple sequence alignments showing the conserved motifs in MpSABATH proteins. The amino-acid numbering corresponds to positions in CbSAMT. SAM binding sites were indicated with red triangles. See Suppl.Fig. S1 for the complete alignment. D, Phylogenetic analysis of SABATH family proteins. MpSABATH proteins were shown in dark blue. See Suppl. Data S1 for the complete phylogram.

Except for Mp*SABATH12*, all the other genes are highly similar in gene structure, each containing two introns in the coding sequence (Fig. 1B). Multiple sequence alignments further showed that the positions of these introns are highly conserved among MpSABATH1-11, while MpSABATH12 is truncated after the first splicing site, likely due to a premature stop codon (Suppl.Fig. S1). The conserved motifs involved in binding the methyl donor SAM (motif A, motif B′ and motif C) (Kato and Mizuno, 2004) were found in all 12 MpSABATH proteins; though MpSABATH8 and MpSABATH9 showed some divergence in the residues that directly bind with SAM in *Clarkia breweri* SAMT (CbSAMT) (Zubieta et al., 2003). In addition, the “YFFF” region downstream of motif C (Kato and Mizuno, 2004), is conserved in MpSABATH1-11 (Fig. 1C; Suppl.Fig. S1). Public transcriptome data retrieved from the MarpolBase Expression database (Kawamura et al., 2022) indicate that all Mp*SABATH* genes are broadly expressed across various tissues, with Mp*SABATH10* showing notably higher expression in dormant spores (Suppl.Fig. S2).

To clarify the evolutionary relationship among SABATH enzymes from *M. polymorpha* and other species, we performed a phylogenetic analysis using sequences from representative species of streptophyte algae and major land plant lineages. Consistent with the previous study (Wang et al., 2024), we observed distinct clustering of algae and land plant proteins, suggesting a monophyletic origin of SABATH family proteins in streptophytes. Though the phylogenetic positions of lycophyte and fern proteins were not conclusive, bryophyte and seed plant proteins segregated into different clades, supporting independent family expansions after the divergence of these plant lineages (Fig. 1D; Suppl.Data S1).

Among the bryophyte SABATHs, proteins from mosses and hornworts each formed a monophyletic clade, with some fern proteins nested in the moss clade. Liverwort SABATH proteins formed two major clades, with one clade containing MpSABATH2 homologs along with some lycophyte proteins, and the other clade comprising all remaining liverwort family members including MpSABATH1, MpSABATH3-12 and the previously reported CsSAMT and CsCAMT from *Conocephalum salebrosum* (Zhang et al., 2019). MpSABATH3/11/12 and MpSABATH10 were positioned close to CsSAMT and CcSAMT, respectively, which might be indicative of their substrate preference (Fig. 1C; Suppl.Data S1).

To explore the biological function of SABATH genes in *M. polymorpha*, we used the CRISPR/Cas9 system to create genome-editing mutants for the full-length genes MpSABATH1-11. Frameshift mutants were obtained for 10 out of the 11 genes except for Mp*SABATH4* (Suppl. Fig. S3). No obvious phenotype was observed in the thallus stage of the mutants except for Mp*sabath2*, which was severely inhibited in thallus growth. Therefore, we focused on Mp*SABATH2* in this study.

### Mp*SABATH2* regulates thallus development and gametangiophore formation

For detailed characterization of Mp*SABATH2* function, we created multiple independent knock-out alleles in different wild-type (WT) accessions. Using the CRISPR/Cas9 nickase system, a large deletion mutant with complete loss of the Mp*SABATH2* genomic locus, Mp*sabath2-1^ld^*, was generated from F1 spore progenies of Tak-1 and Tak-2. Three other genome-editing alleles, Mp*sabath2-2^ge^*♂, Mp*sabath2-3^ge^*♀, and Mp*sabath2-4^ge^*♂ were created using CRISPR/Cas9 with a single gRNA from the spore progenies of recombinant inbred lines (Tomizawa et al., 2023). Mp*sabath2-2^ge^*♂ was backcrossed with the female parent (Rit-2) to remove the CRISPR T-DNA, which generated the Mp*sabath2-2^ge^*♀ female segregant (Suppl. Fig. S4).

All the Mp*sabath2* mutant alleles showed consistent phenotypes. Under regular continuous white light growth conditions (cW), the mutants developed very small, dark green thalli that were curled upwards, and no gemma cups were formed (Fig. 2A-C; Suppl. Fig. S5A-D,G-H; S8A-B). In line with the dark-green color, chlorophyll measurements revealed that both chlorophyll *a* and *b* were significantly accumulated in the mutants compared to WT (Fig. 2D; Suppl. Fig. S5E-F,I-J). Meanwhile, cross sections of Mp*sabath2* thalli showed no substantial reduction in cell sizes compared to WT (Suppl. Fig. S6), suggesting that the smaller plant size is more likely attributable to reduced cell proliferation rather than defective cell expansion.

**Figure 2.**
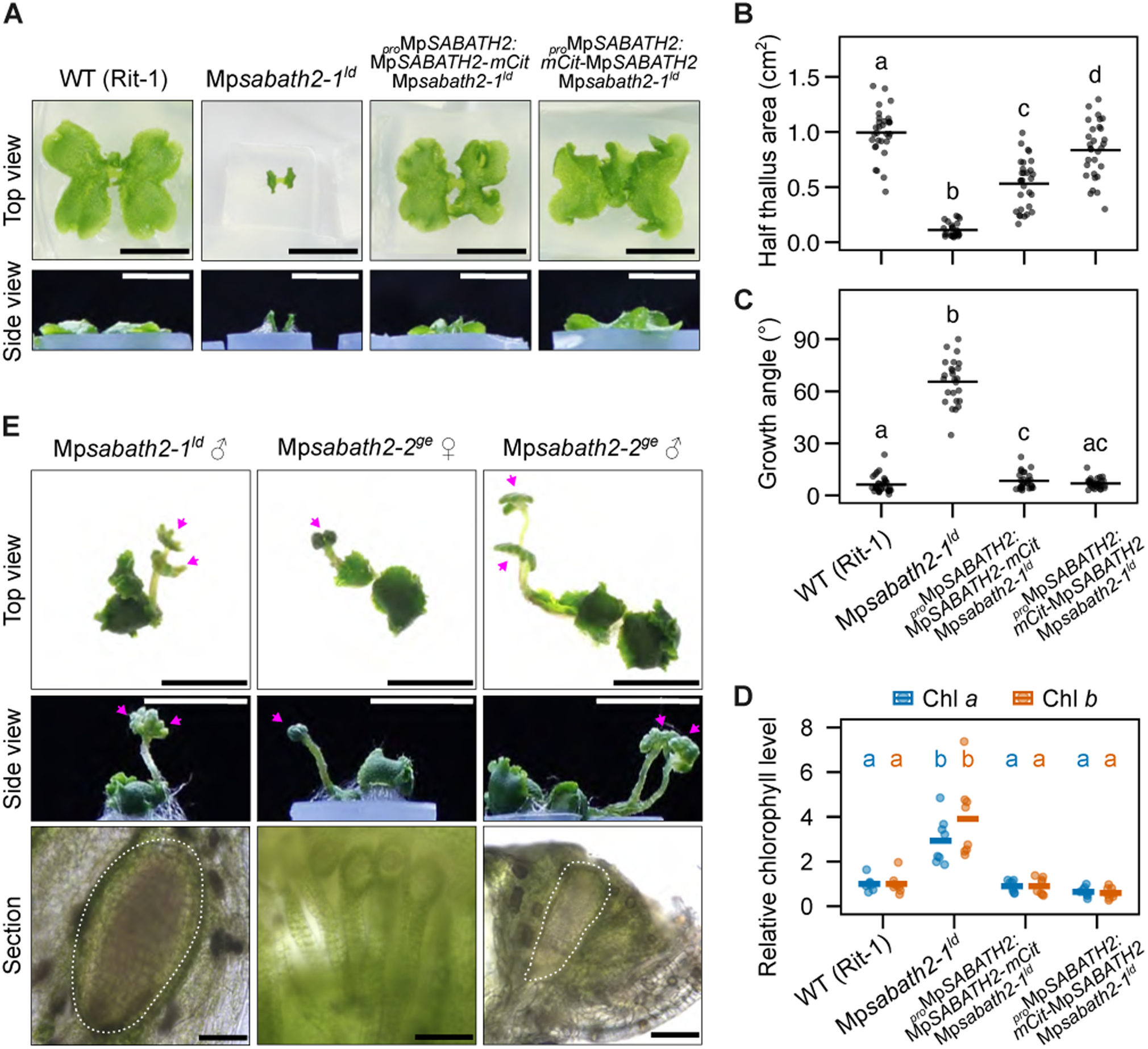
Loss of Mp*SABATH2* affected thallus morphology and promoted sexual reproduction. A, Photos of 14-day-old plants of wild-type (WT), Mp*sabath2-1^ld^* and the complementation lines grown from gemmae under continuous white light (cW) conditions. Bars = 1 cm. B-D, Measurements of thallus area (B), growth angle (C), chlorophyll *a* and chlorophyll *b* (D) from plants shown in (A). Non-overlapping letter combinations indicate significant statistical difference (*P* < 0.05). For B-C, pairwise Mann-Whitney *U* test with Benjamini-Hochberg (B-H) correction was used. For D, Tukey-Kramer test was used. E, Photos of 39-day-old Mp*sabath2* plants grown under cW, showing sexual reproduction in both male and female mutants. Magenta arrowheads in the camera photos indicate gametangiophores. White dashed lines in the longitudinal sections indicate antheridia. Bars = 1 cm (top view and side view photos); Bars = 100 µm (sections).

In addition, both male and female Mp*sabath2* mutants formed sexual reproductive branches, i.e. gametangiophores, when cultured for 39 days under cW, despite the absence of far-red light (FR) enrichment required for sexual reproduction in WT plants (Chiyoda et al., 2008; Inoue et al., 2019) (Fig. 2E). Longitudinal sections confirmed that antheridia and archegonia were properly formed in these gametangiophores (Fig. 2E).

To further investigate the influence of Mp*sabath2* on sexual reproduction, we observed the mutants under the FR-enriched condition (cW+FR), which induces sexual reproduction along with apical meristem dormancy in *M. polymorpha* (Inoue et al., 2019; Streubel et al., 2023). Consistent with the promotion of sexual reproduction under cW, the time to form the first visible gametangiophore was significantly shorter in the Mp*sabath2* mutants (9.7±0.5, 12.1±1.4, 9.3±1.7) than that of male (13.2±1.2) and female (17.9±1.7) WT plants under cW+FR (Suppl. Fig. S7A-B). Moreover, as the gametangiophores in Mp*sabath2* mutants formed before any branching occurred, the thallus branching was entirely terminated. No new branches were formed even after 21 days of FR irradiation, which further suppressed thallus expansion in these plants (Suppl. Fig. S7A). We crossed male and female Mp*sabath2* mutants with each other and WT partners. Sporangia and mature spores were formed from all combinations of crossings, indicating that Mp*SABATH2* loss-of-function does not affect fertility (Suppl. Fig. S7C).

To confirm that Mp*SABATH2* is the causal gene for the observed phenotypes, we complemented the Mp*sabath2-1^ld^* mutant by two constructs expressing Mp*SABATH2* under its own promoter, with the monomeric Citrine fused to either the amino- or carboxyl-terminus of the coding region. Both complementation lines restored all the Mp*sabath2^ld^* phenotypes to the level of WT plants, including thallus size (Fig. 2A-B), thallus growth angle (Fig. 2A, C), chlorophyll accumulation (Fig. 2D) and the formation of gemmae and gemma cups (Suppl. Fig. S7A). Spontaneous sexual reproduction under cW was suppressed in the complementation lines, and the timing of gametangiophore formation under cW+FR was restored (Suppl. Fig. S7A-B).

### Mp*sabath2* transcriptome resembled features of FR response

Mp*SABATH2* loss-of-function caused upward growth of the thallus and activated sexual reproduction, which is typical of FR response in *M. polymorpha* (Fredericq, 1964; Inoue et al., 2019). To examine if such phenotypic similarity is supported by gene expression changes, we compared the transcriptomes of male and female Mp*sabath2-2^ge^* mutants with WT plants under cW conditions, as well as the transcriptomes of WT plants exposed to different light conditions.

A wide range of gene expression changes was observed in 14-day-old Mp*sabath2-2^ge^* mutants. Both male and female mutants showed more than 1,800 up- and nearly 1,000 down-regulated genes, with 1,484 up- and 785 down-regulated genes shared between the two sexes. Among these, around 1/3 of the up- or down-regulated genes were also significantly up- or down-regulated in response to FR treatment in WT plants (Fig. 3A). Furthermore, when we performed a correlation analysis among these datasets, Pearson’s correlation coefficients exceeded 0.7 for three of the four comparisons between Mp*sabath2* mutants and WT FR responses, indicating a strong positive correlation between Mp*sabath2* and FR-influenced genes (Fig. 3B).

**Figure 3.**
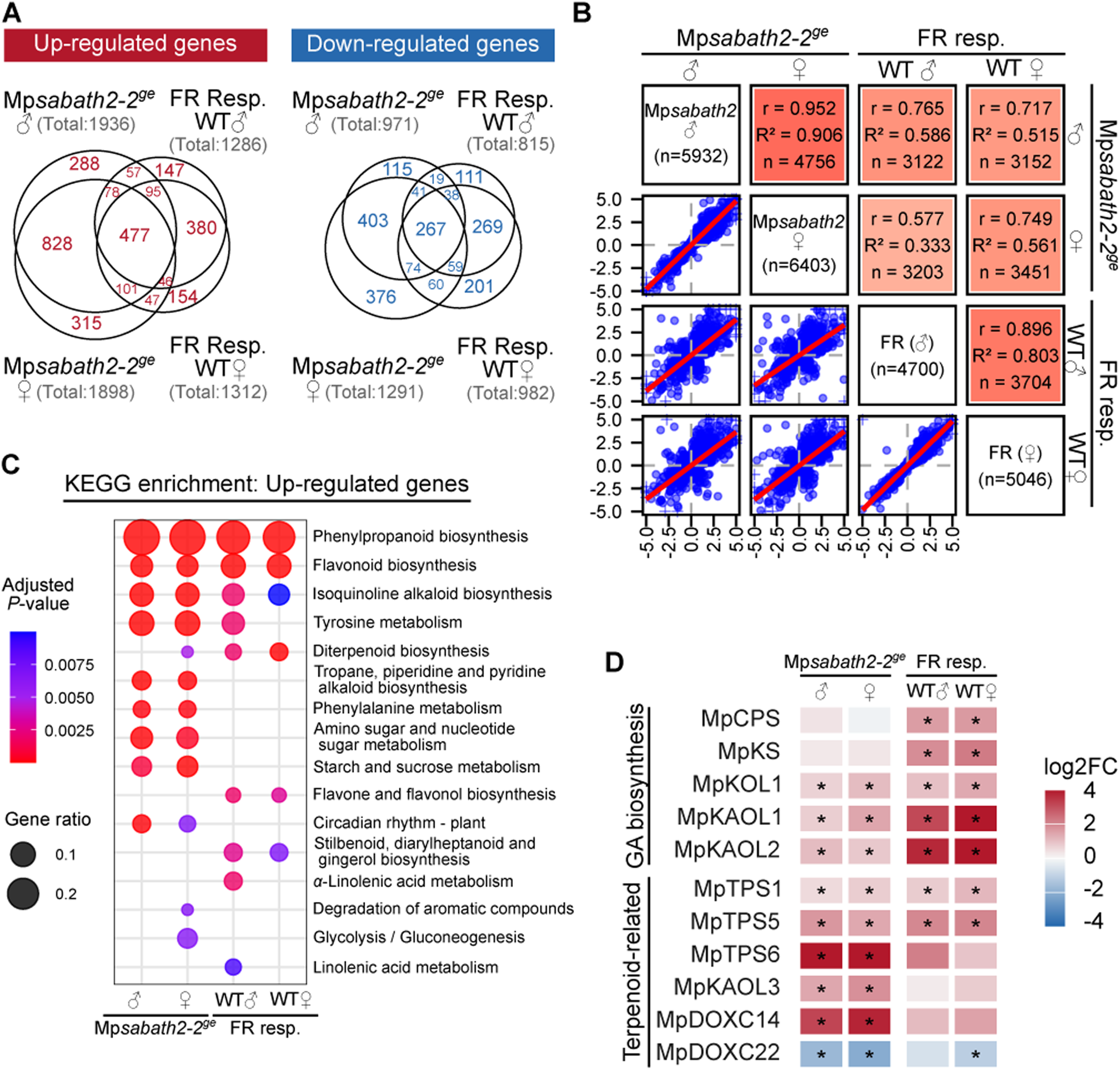
Gene expression changes in Mp*sabath2* and far-red light (FR)-iraddiated WT plants. A, Euler diagram showing differentially expressed genes (DEGs) in Mp*sabath2-2^ge^*mutants grown under cW and DEGs between cW+FR and cW conditions in WT plants (“FR Resp.”). DEGs were determined by DESeq2 with an adjusted *P*<0.01 and |log2(Fold Change)|>0.585). B, Pairwise correlation analysis of gene expression changes. Upper-right triangle matrix: pairwise correlation of different datasets. The numbers are the Pearson correlation coefficients (r), the coefficient of determination (R^2^) and the sample size (n). The color intensity indicates the degree of pairwise correlation. Lower-left triangle matrix: scatter plot showing z-scores calculated from log2 fold changes of each gene. C, Enrichment analysis of KEGG (Kyoto Encyclopedia of Genes and Genomes) pathways in genes up-regulated in Mp*sabath2^ge^* or upon FR irradiation. D, Heatmap showing DEGs that are putatively involved in GA biosynthesis or terpenoid metabolism.

As many SABATH family enzymes are involved in primary or secondary metabolism, KEGG pathway enrichment analysis was performed to identify the metabolic pathways influenced by Mp*sabath2* (Fig. 3C). While no plant-related pathway was significantly enriched for down-regulated genes in Mp*sabath2-2^ge^* (Suppl. Fig. S8), up-regulated genes in Mp*sabath2-2^ge^*were most enriched in phenylpropanoid and flavonoid biosynthesis, isoquinoline, alkaloid biosynthesis and tyrosine metabolism. These pathways were also top-ranked among FR-induced genes, further supporting that Mp*SABATH2* loss-of-function evoked transcriptional changes similar to those of the FR response. Diterpenoid biosynthesis is a pathway up-regulated in the FR response but not clearly enriched in Mp*sabath2* up-regulated genes, so we further checked the expression patterns of related genes (Fig. 3D). FR treatment significantly increased the expression of Mp*TPS1*, Mp*TPS5* and several genes involved in gibberellin biosynthesis (Sun et al., 2023). While some of these genes were also up-regulated in Mp*sabath2-2^ge^*, Mp*sabath2-2^ge^* did not significantly enhance the expression of Mp*CPS* or Mp*KS*, but instead strongly induced the expression of Mp*TPS6*, Mp*KAOL3* and Mp*DOXC14*, which are tentatively associated with different diterpenoid metabolites, such as *cis*-kolavenol (Kumar et al., 2016).

Mp*sabath2-2^ge^* also affected metabolic pathways that did not respond to FR treatment. In particular, the starch and sucrose metabolism was enriched in Mp*sabath2-2^ge^* up-regulated genes. We examined the starch levels in Mp*sabath2-2^ge^* with the iodine–starch test. Compared with WT, Mp*sabath2-2^ge^* showed strong starch accumulation across the thallus, most prominent in the apical region (Fig. 4A). Consistently, we observed starch granules more frequently in the chloroplasts of Mp*sabath2-2^ge^*, as shown in the assimilation filaments near the apical region (Fig. 4B), or ventral epidermal cells (Suppl. Fig. S6B).

**Figure 4.**
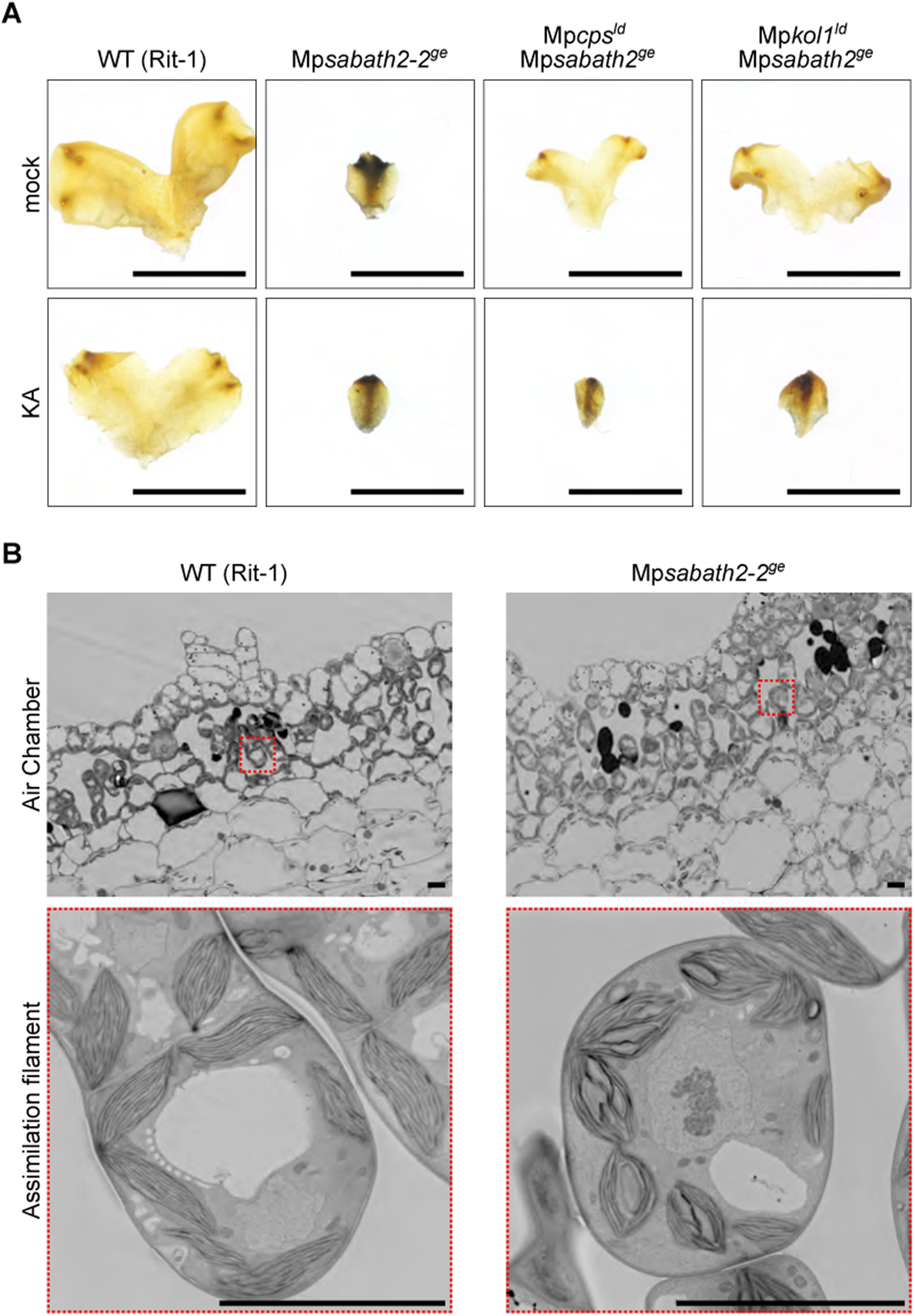
Mp*sabath2* mutation caused starch accumulation. A, Iodine staining showing different levels of starch accumulation in 14-day-old plants grown under cW. Bars = 1 cm. B, Ultra-thin sections near the apical region, showing the formation of starch granules in chloroplasts of Mp*sabath2-2^ge^* plants. Images were taken with the field-emission scanning electron microscope (FE-SEM). Bars = 10 µm.

### Inhibition of gibberellin precursor biosynthesis suppressed Mp*sabath2* phenotypes

SABATH family enzymes are known to be involved in the metabolism of many plant hormones, including auxin, GA, JA, and SA. Recent research has shown that in *M. polymorpha*, although the canonical GAs are absent, the biosynthesis of GA precursors such as *ent*-kaurenoic acid (KA) is required for proper FR responses (Sun et al., 2023). Given the similarity of Mp*SABATH2* loss-of-function and FR response, we investigated the genetic interaction between Mp*SABATH2* and the GA-related metabolic pathway in *M. polymorpha*.

First, we tested the effect of paclobutrazol (PAC), a cytochrome P450 (CYP) enzyme inhibitor that mainly targets the kaurene oxidase (KO) in GA biosynthesis (Rademacher, 2000). Treatment with 20 µM of PAC in the medium largely suppressed Mp*sabath2* phenotypes: the thallus size was significantly increased, and the gemma and gemma cup formation was restored in the Mp*sabath2-2^ge^* mutants (Suppl.Fig. S9A-C). Mp*sabath2* gemmae formed in the presence of PAC were morphologically indistinguishable from those of WT, which allowed us to grow Mp*sabath2* mutants from gemmae in all experiments (Suppl.Fig. S9A). Nevertheless, such phenotype suppression by PAC treatment was also incomplete. The thallus sizes of PAC-treated Mp*sabath2-2^ge^*were much smaller than those of WT plants, and the thallus growth angles were even higher in PAC-treated plants than the mock group.

Next, we genetically disrupted GA precursor biosynthesis by constructing double mutants of Mp*sabath2-2^ge^*with either Mp*cps^ld^* or Mp*kol1^ld^*. Both enzymes act upstream of KA, and the mutations suppressed Mpsabath2 phenotypes in a way similar to PAC treatment. The thallus sizes were partially rescued (Fig. 5A,C), the gemma and gemma cup formation was restored, and the spontaneous sexual reproduction was suppressed (Fig. 5B). Mp*cps^ld^* and Mp*kol1^ld^* mutations also suppressed starch accumulation in Mp*sabath2-2^ge^*mutants (Fig. 4A). Furthermore, application of 2 µM KA in the medium reverted all phenotypes to the levels of Mp*sabath2-2^ge^* (Figs. 5A-C, 4A), which confirmed the involvement of GA metabolism in Mp*sabath2*. Similar to the situation in PAC treatment, the thallus growth angle did not significantly change in the double mutants, and only increased in the Mp*kol1^ld^*Mp*sabath2-2^ge^* line in response to KA treatment (Fig. 5D).

**Figure 5.**
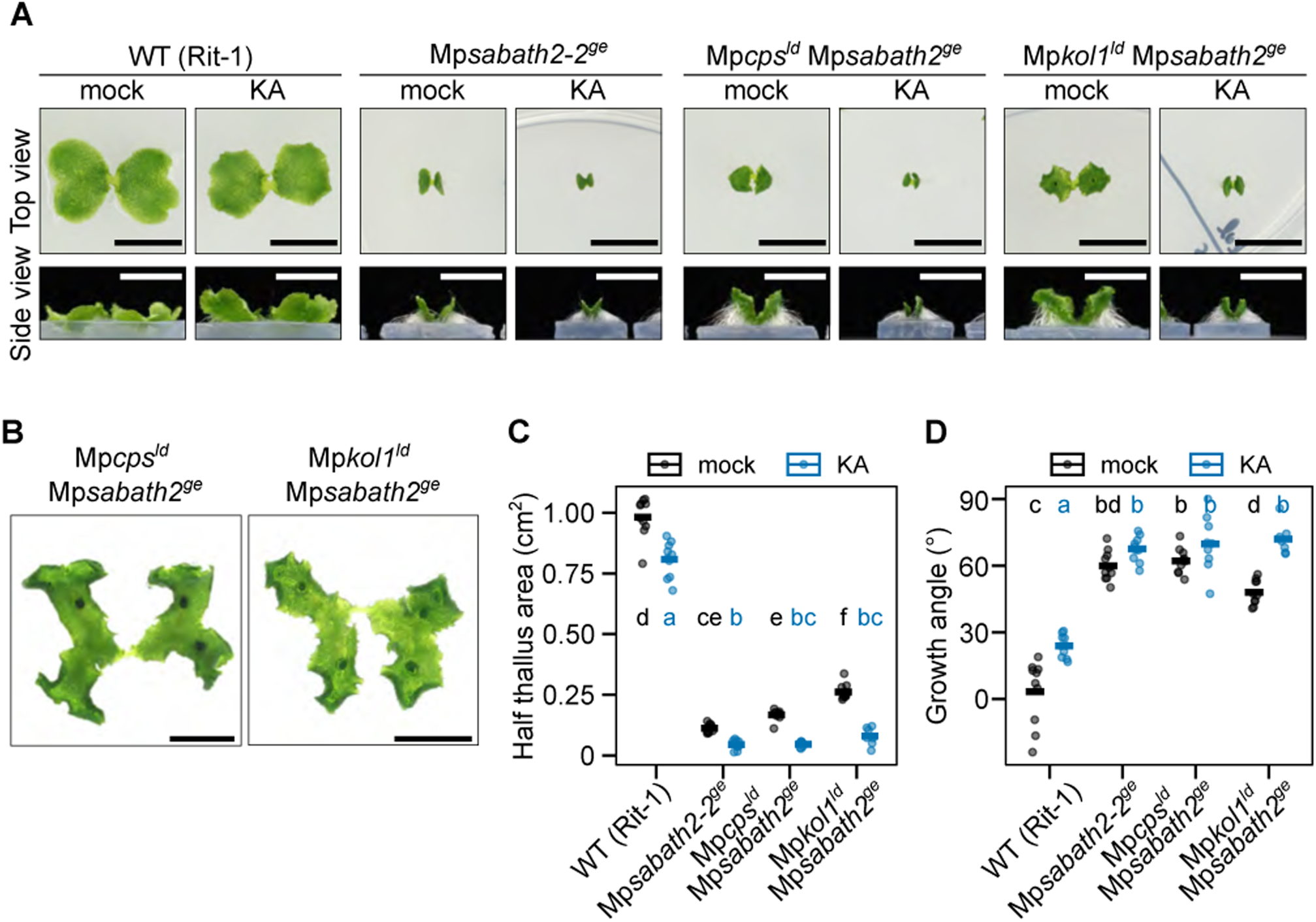
KA biosynthesis deficiency suppressed Mp*sabath2* phenotypes. A, Photos of 14-day-old WT, Mp*sabath2^ge^*, Mp*cps^ld^*Mp*sabath2^ge^* and Mp*kol1^ld^* Mp*sabath2^ge^*plants grown from gemmae with or without 2 μM KA under cW conditions. Bars = 1 cm. B, Photos of 29-day-old Mp*cps^ld^* Mp*sabath2^ge^* and Mp*kol1^ld^* Mp*sabath2^ge^* plants grown under cW conditions, showing the formation of gemma cups. Bars = 1 cm. C-D, Measurements of thallus area (C) and growth angle (D) from plants shown in (A). Non-overlapping letter combinations indicate significant statistical difference based on the Tukey-Kramer test (*P* < 0.05).

## Discussion

The SABATH family methyltransferases catalyze the methylation of many plant-specialized metabolites, including several plant hormones. The emergence of this plant-specific family can be traced back to streptophyte algae. Phylogenetic analyses suggested that it arose from a few, or a single ancestral gene in the common ancestor of land plants, and later underwent independent expansion in different lineages (Wang et al., 2024) (Fig. 1C). It is common for SABATH enzymes of distinct evolutionary origins to acquire similar enzymatic activities. BAMT, SAMT, or bifunctional benzoic/salicylic acid MTs (BSATs) have been reported from angiosperm, gymnosperm, and liverwort species (Ross et al., 1999; Chen et al., 2003; Chaiprasongsuk et al., 2018; Zhang et al., 2019), which evolved independently in their own lineage. Similarly, JMTs of angiosperms and gymnosperms belong to different ortholog groups (Chaiprasongsuk et al., 2018). Our investigation agrees with the previous analyses that all liverwort SABATHs belong to one or two monophyletic clades well-separated from angiosperm and gymnosperm proteins, and the substrate preference likely evolved independently in this plant lineage.

Meanwhile, IAMTs and GAMTs are known to form monophyletic clades that diverged early from other seed plant SABATHs (Chaiprasongsuk et al., 2018; Zhang et al., 2020; Wang et al., 2024), which possibly reflects their conserved function in hormone deactivation and growth regulation. We observed a similar pattern for MpSABATH2 orthologs, which might indicate a unique functional specialization in the liverwort lineage.

Our results showed that Mp*SABATH2* is a key regulator of plant development, affecting multiple processes including growth, reproduction, photosynthesis, and starch metabolism. Loss of Mp*SABATH2* function strongly inhibited thallus growth and increased the growth angle. Also, the gene is required for maintaining asexual reproduction, i.e. gemma and gemma cup formation, while suppressing sexual reproduction under regular light conditions (Fig. 2). To our knowledge, these developmental defects were more severe than those caused by loss-of-function or gain-of-function of SABATH enzymes in other species (Seo et al., 2001; Qin et al., 2005; Varbanova et al., 2007). This leads to the hypothesis that MpSABATH2 might be involved in plant hormone metabolism, serving as a deactivation enzyme or biosynthetic gene. However, the MpSABATH2 homolog in the liverwort *C. salebrosum*, CsSABATH1, showed no activity towards IAA, GA, JA, SA, or ABA in a previous study (Zhang et al., 2019), suggesting that the substrate might be different from common hormones active in seed plants.

Some of the phenotypes of Mp*sabath2* mutants resemble the responses to far-red enriched light conditions in WT plants, such as the change in growth angle and transition to sexual reproduction (Fredericq and de Greef, 1966; Inoue et al., 2019). This was further supported by our transcriptome analysis (Fig. 3). One possibility is that MpSABATH2 alters the metabolism of a hormone that bypasses the environmental clues to trigger these responses. The GA-related pathway is required for FR response in *M. polymorpha*, and GA biosynthetic mutants caused phenotypes opposite to those of Mp*sabath2*. Mp*cps* or Mp*kol1* mutations increased thallus size and delayed sexual reproduction under cW+FR conditions (Sun et al., 2023), suggesting that GA-related compounds might be candidate substrates for MpSABATH2. Our chemical treatments and genetic analyses partially supported this idea. PAC treatment, or mutations in Mp*CPS or* Mp*KOL1* partially rescued phenotypes of Mp*sabath2*, which was restored by KA treatment (Fig. 5). In fact, Mp*cps* Mp*sabath2* and Mp*kol1* Mp*sabath2* showed much higher sensitivity to KA treatment under cW than that observed for Mp*cps* or Mp*kol1* single mutants in the previous research (Sun et al., 2023), which can be interpreted as a result of defective GA deactivation in Mp*sabath2*. However, as the bioactive form of GA-related hormone is still unclear in *M. polymorpha*, we do not have any direct evidence for this hypothesis. Given the up-regulated expression of GA biosynthesis genes in Mp*sabath2* (Fig, 3D), it remains possible that GA homeostasis is indirectly altered as well.

On the other hand, blocking GA biosynthesis did not fully rescue the phenotypes of Mp*sabath2*, especially the plant size and growth angle (Fig.5), suggesting the involvement of other growth regulating mechanisms. One candidate is the cytokinin pathway, as deficiency in cytokinin signaling or biosynthesis in *M. polymorpha* inhibits thallus growth and gemma cups formation, as well as increases growth angle (Aki et al., 2019; Komatsu et al., 2025). There is no report of plant-sourced methylated cytokinin yet, although mono- and dimethylated *N*^6^-(Δ^2^-isopentenyl)adenine produced by the bacteria *Rhodococcus fascians* showed cytokinin activity in Arabidopsis (Radhika et al., 2015). The functional relevance of MpSABATH2 to cytokinin metabolism remains to be examined.

Overall, our findings identified a novel SABATH enzyme that is critical for plant development in the liverwort *M. polymorpha*. Characterizing its biochemical function will be an important next step toward deciphering the relevant endogenous growth-regulating compounds and how these metabolic pathways evolved across land plants.

## Materials and Methods

### Plant materials and growth conditions

Wild-type *Marchantia polymorpha* Takaragaike-1 (Tak-1), Takaragaike-2 (Tak-2) (Ishizaki et al., 2008) and recombinant inbred lines of Takaragaike (Rit-1 and Rit-2) (Tomizawa et al., 2023) were used in this study.

Unless otherwise specified, *M. polymorpha* plants were aseptically cultured on half-strength Gamborg’s B5 media under continuous white light (cW, 40-50 µmol m^-2^ s^-1^) from white fluorescent lamps (CCFLs, OPT-40C-N-L, Optrom, Japan) at 22°C.

### Plasmid construction

Construction of the vectors for CRISPR-mediated genome editing with single guide RNAs (gRNAs) was performed according to (Sugano et al., 2014). Target sites for genome editing were selected with the CRISPRdirect website (Naito et al., 2015). Annealed oligos were inserted into the BsaI site of pMpGE_En03 and then transferred to the binary vector pMpGE011 using Gateway LR Clonase II Enzyme Mix (Thermo Fischer Scientific) according to the manufacturer’s instructions.

To generate large-deletion mutants of the entire gene locus, a nickase-type CRISPR/Cas9 genome editing system was employed (Koide et al., 2020). Two pairs of gRNAs were designed targeting sequences upstream and downstream of Mp*SABATH2*, Mp*CPS* or Mp*KOL1*. Annealed oligonucleotide duplexes were cloned separately into the BsaI sites of pMpGE_En04, pBC-GE12, pBC-GE23, and pBC-GE34. The gRNA expression cassettes from the three pBC-GE vectors were subsequently integrated into the BglI site of the pMpGE_En04 vector. The resulting entry clones were transferred into the pMpGE_017 or pMpGE018 destination vector using Gateway LR Clonase II (Thermo Fisher Scientific).

For complementation of Mp*sabath2-1^ld^*, a genomic fragment spanning 5 kb upstream of the Mp*SABATH2* promoter to 1 kb downstream of the stop codon was amplified by PCR using primers MpSABATH2_genome_primer1_CACC and MpSABATH2_genome_primer2, and cloned into pENTR-dTOPO. mCitrine with a GGSG linker sequence was introduced at either the N-terminus or C-terminus using the In-Fusion HD cloning kit (TaKaRa). The resulting pENTR-dTOPO-MpSABATH2-mCitrine and pENTR-dTOPO-mCitrine-MpSABATH2 constructs were transferred into pMpGWB101 (Ishizaki et al., 2015) using Gateway LR Clonase II.

The oligonucleotides used in this study are listed in Suppl. Table S2.

### Transformation of *M. polymorpha*

Transformation of *M. polymorpha* was carried out following procedures described in previous publications (Ishizaki et al., 2008; Kubota et al., 2013). For transformation using regenerating thalli, apical regions were removed from 10∼12-day-old thalli, and the basal fragments were cultured on ½ Gamborg’s B5 medium containing 1.0% (w/v) sucrose and 1.0% (w/v) agar for three days under continuous white light for regeneration. For sporeling transformation, spores were sterilized and cultured in the liquid 0M51C medium with containing 2% (w/v) sucrose for 7-10 days. The regenerating thalli or sporelings were co-cultured with the Agrobacterium strain GV2260 harbouring the binary vector in the liquid 0M51C medium containing 2% (w/v) sucrose and 100 μM acetosyringone under continuous white light with agitation at 130 rpm for three days. Transformants were washed and selected on ½ Gamborg’s B5 medium plates containing 100 mg L^−1^ cefotaxime and proper antibiotics.

### Generation of mutants

For initial screening, Mp*sabath* mutants were generated from Tak-1 plants with thallus transformation (Kubota et al., 2013). The Mp*sabath2-1^ld^* mutant was generated from the F1 progeny of Tak-1 and Tak-2 with sporeling transformation. The mutants Mp*sabath2-2^ge^* ♂, Mp*sabath2-3^ge^* ♀ and Mp*sabath2-4^ge^* ♂ were generated from the F1 progeny of Rit-1 and Rit-2 with sporeling transformation. Mp*sabath2-2^ge^* ♂ was then backcrossed with Rit-2 to remove the CRISPR vector, which generated the female segregate Mp*sabath2-2^ge^* ♀. Complementation lines of Mp*sabath2-1^ld^*were generated with thallus transformation.

As for the double mutants, Mp*cps^ld^* Mp*sabath2^ge^*was generated from Mp*sabath2-2^ge^* ♂ by thallus transformation using pMpGE017-MpCPS-LD from (Sun et al., 2023). For Mp*kol1^ld^* Mp*sabath2^ge^*, a Mp*kol1^ld^*mutant with large deletion was first generated from the F1 progeny of Tak-1 and Tak-2 with sporeling transformation, then Mp*SABATH2* was mutated via thallus transformation.

### Morphological observation

Thallus sizes and growth angles were measured from half thallus pieces of 14-day-old plants following the method of (Sun et al., 2023).

Sexual reproduction was induced by supplementing cW with far-red light (cW+FR, 40-50 µmol m^-2^ s^-1^). The number of days required for gametangiophore induction was counted from the start of far-red light supplementation until gametangiophores became visible at the thallus apex by naked-eye observation.

### Chlorophyll measurement

Plants grown for 14 days under continuous white light after gemma germination were weighed, and chlorophyll was extracted with *N*,*N*-dimethylformamide. Absorbance at 663.8 nm and 646.8 nm was measured using the Powerscan4 plate-reader for chlorophyll *a* and chlorophyll *b*, respectively. The relative chlorophyll contents were calculated in reference to the control samples.

### Iodine staining of starch

The iodine staining of starch for *M. polymorpha* was performed following the methods of (Hostettler et al., 2011). Plants grown for 14 days under cW were bleached in 80% (v/v) ethanol containing 0.1% (v/v) 2-mercaptoethanol at 80°C and stained with Lugol solution [0.34% (w/v) I₂, 0.68% (w/v) KI] for 5 minutes. After staining, the samples were washed in water for three times, 2 minutes for each time.

### Sectioning and FE-SEM observation

To prepare ultrathin sections, 14-day-old thalli were fixed overnight at 4°C in 2.5% (v/v) glutaraldehyde and 2% (w/v) paraformaldehyde in 50 mM phosphate buffer (pH 7.2). After washing with 50 mM phosphate buffer, samples were post-fixed with 2% (w/v) osmium tetroxide for 2 hours at room temperature, dehydrated through a graded ethanol series [25%, 50%, 80%, 99%, 100% (v/v)], and embedded in epoxy resin. Ultrathin sections (150 nm thick) were cut with a diamond knife using an ultramicrotome (Ultracut UCT; Leica, Wetzlar, Germany), mounted on silicon wafers (380-μm thick; Canosis, Tokyo, Japan), and stained with 2% (w/v) uranyl acetate and lead citrate. Samples were observed with field emission scanning electron microscopes (JSM-7900F; JEOL, Tokyo, Japan).

### RNA-sequencing

For RNA-sequencing, Mp*sabath2-2^ge^* ♂ ♀ mutants and the WT Rit-1 and Rit-2 plants were cultured from gemmae for 14 days under cW. In addition, Rit-1 and Rit-2 plants were cultured for 3 days under cW+FR following 11 days under cW to evaluate the response to FR. Considering the difference in plant size, the apical region of WT plants or whole thalli of Mp*sabath2-2^ge^* were collected, flash-frozen with liquid nitrogen and stored at -80 °C until RNA extraction.

RNA extraction was performed using the RNeasy Plant Mini Kit (QIAGEN) following the manufacturer’s protocol. Quality control was performed using a Bioanalyzer 2100 (Agilent) with the RNA6000 Pico Kit (Agilent). mRNA was enriched using the NEBNext Poly(A) mRNA Magnetic Isolation Module (New England Biolabs, #E7490), and libraries were prepared using the NEBNext Ultra II Directional RNA Library Prep Kit for Illumina (New England Biolabs, #E7760). Libraries were amplified using NEBNext Multiplex Oligos for Illumina (96 Unique Dual Index Primer Pairs Set 2, New England Biolabs, #E6442). Quality control was performed using the Bioanalyzer High Sensitivity DNA assay (Agilent), and sequencing was performed on a NextSeq 500 using the NextSeq 500/550 High Output Kit v2.5 (75 cycles) (Illumina). Samples were de-multiplexed using BaseSpace Sequence Hub (Illumina).

For RNA-seq data analysis, the reference *M. polymorpha* MpTak_v7.1 (U+V) genome, gene structure annotation, and KEGG annotation were obtained from Marpolbase (Tanizawa et al. 2025). The GFF annotation file was converted to GTF format using the agat_convert_sp_gff2gtf.pl script from AGAT (version 0.8.0) (Dainat et al., 2025). Sequencing reads were processed through the nf-core/rnaseq pipeline (version 3.13.2) (Patel et al., 2025), where quality trimming of raw reads was performed by fastp (Chen et al., 2018). Transcript abundance was quantified using Salmon (Patro et al., 2017) in pseudo-alignment mode. Salmon quantification included bias correction parameters --seqBias --gcBias to account for sequence-specific and GC content biases. Differential gene expression analysis was performed using the DESeq2 package (Love et al., 2014). Count matrices were imported using tximport (Soneson et al., 2016) to aggregate transcript-level abundances to gene-level counts. Sample metadata was constructed based on experimental factors including light conditions and genotype. DESeq2 analysis was conducted for each comparison with appropriate design formulas and reference levels using custom wrapper functions. Genes were designated as differentially expressed when they met the criteria of adjusted p-value < 0.01 and absolute log2FC > 1 (alternatively, > 0.585 for Euler diagram). To identify shared and condition-specific gene expression patterns across experimental conditions, log2 fold changes were standardized using z-score normalization, and Pearson correlation coefficients were calculated between pairwise comparisons. Overlapping differentially expressed genes among conditions were visualized using Euler diagrams created with the eulerr package. KEGG pathway enrichment analysis was conducted using the clusterProfiler package (Yu et al., 2012). Statistical significance was evaluated through hypergeometric testing followed by Benjamini-Hochberg multiple testing correction, with pathways showing adjusted p-value < 0.01 considered as significantly enriched. Expression patterns of selected gene sets were displayed as heatmaps showing log2 fold changes across multiple comparisons. All data visualizations were created using ggplot2 and related R packages with assistance from generative AI (Claude Sonnet 4).

### Phylogenetic analysis

To conduct phylogenetic analysis of the SABATH protein family, we performed a blastp search (version 2.12.0) using the MpSABATH2 amino acid sequence from MpTak-1 v7.1 as the reference query. This search was conducted against annotated protein sequences from published genomes (see Suppl. Table S3 for the genome sources) and the OneKP database entry ILBQ for *Conocephalum salebrosum* (chemotype of *Conocephalum conicum*). We retrieved blast results with E-values below 1e-10 using seqkit software (version 2.9.0). Additionally, ten SABATH proteins (SABATH1 through SABATH10) from *Picea abies* were incorporated manually into the dataset. We performed sequence alignment using MAFFT (version 7.526) with the FFT-NS-i algorithm, then refined the alignment by eliminating positions containing more than 80% gaps using trimAl (version 1.4. rev15). For phylogenetic tree construction, we employed RAxML-NG (version 1.2.1) with 1000 bootstrap replicates and the LG+G4m substitution model, which was selected based on ModelTest-NG (version 0.1.7) recommendations. The resulting phylogenetic tree was visualized using the ggtree package in R.

### Data Availability Statement

The RNA-seq data underlying this article are available in DNA Data Bank of Japan (DDBJ) at www.ddbj.nig.ac.jp, and can be accessed with PRJDB38067.

## Supporting information

Supplemental FiguresTables

## Acknowledgments

We thank Ryunosuke Kusunoki (Kyoto University) for the construction of Mp*kol1^ld^* mutant; Takao Koeduka and Kenji Matsui (Yamaguchi University), Maiko Okabe and Takuya Segawa (Kyoto University) for their contributions to preliminary data not shown in this manuscript; and Feng Chen (University of Tennessee) for comments and advice. We also thank Takehumi Kondo and Yukari Sando (NGS core facility of the Grad. Sch. of Bio., Kyoto Univ.) for supporting the RNA-seq analysis. Computation time was provided by the Supercomputer System, Institute for Chemical Research, Kyoto University.

## Author Contributions

S.K., T.U., M.K., and T.Ko. planned and conceived the project. S.K., E.S., M.It, I.N., T.Ka., M.Iw., Y.Y. performed experiments. S.K., E.S., T.Ka, M.Iw, R.S., Y.Y., S. Yamao., S. Yamag., T.U., M.K. and T.Ka. analyzed the data. S.K., E.S., R.S., Y.Y. S.Yamao., and T.Ko wrote the manuscript with input from all authors.

## References

1. Abbas, M., Hernández-García, J., Blanco-Touriñán, N., Aliaga, N., Minguet, E.G., Alabadí, D., et al. (2018) Reduction of indole-3-acetic acid methyltransferase activity compensates for high-temperature male sterility in Arabidopsis. Plant Biotechnology Journal. 16: 272–279.

2. Aki, S.S., Mikami, T., Naramoto, S., Nishihama, R., Ishizaki, K., Kojima, M., et al. (2019) Cytokinin signaling is essential for organ formation in *Marchantia polymorpha*. Plant and Cell Physiology. 60: 1842–1854.

3. Azar, M., Goldbecker, E., Karpovsky, D., Shpilman, M., Breker, M., de Vries, J., et al. (2025) Shared abscisic acid biosynthesis pathway across 600 million years of streptophyte evolution. Plant Physiol. 198: kiaf121.

4. Chaiprasongsuk, M., Zhang, C., Qian, P., Chen, X., Li, G., Trigiano, R.N., et al. (2018) Biochemical characterization in Norway spruce (*Picea abies*) of SABATH methyltransferases that methylate phytohormones. Phytochemistry. 149: 146–154.

5. Chen, F., D’Auria, J.C., Tholl, D., Ross, J.R., Gershenzon, J., Noel, J.P., et al. (2003) An *Arabidopsis thaliana* gene for methylsalicylate biosynthesis, identified by a biochemical genomics approach, has a role in defense. The Plant Journal. 36: 577–588.

6. Chen, S., Zhou, Y., Chen, Y., and Gu, J. (2018) fastp: an ultra-fast all-in-one FASTQ preprocessor. Bioinformatics. 34: i884–i890.

7. Chini, A., Monte, I., Zamarreño, A.M., García-Mina, J.M., and Solano, R. (2023) Evolution of the jasmonate ligands and their biosynthetic pathways. New Phytologist. 238: 2236–2246.

8. Chiyoda, S., Ishizaki, K., Kataoka, H., Yamato, K.T., and Kohchi, T. (2008) Direct transformation of the liverwort *Marchantia polymorpha* L. by particle bombardment using immature thalli developing from spores. Plant Cell Reports. 27: 1467–1473.

9. Dainat, J., Cannoodt, R., Soares, A., Ruano, D.G., Hereñú, D., Murray, D.K.D., et al. (2025) AGAT: Another Gff Analysis Toolkit to handle annotations in any GTF/GFF format. (Version v0.8.0). Zenodo. https://zenodo.org/records/16317950.

10. D’Auria, J.C., Chen, F., and Eran Pichersky (2003) The SABATH family of MTS in *Arabidopsis thaliana* and other plant species. In Recent Advances in Phytochemistry. pp. 253–283 Elsevier.

11. Denoeud, F., Carretero-Paulet, L., Dereeper, A., Droc, G., Guyot, R., Pietrella, M., et al. (2014) The coffee genome provides insight into the convergent evolution of caffeine biosynthesis. Science. 345: 1181–1184.

12. Farmer, E.E., and Ryan, C.A. (1990) Interplant communication: airborne methyl jasmonate induces synthesis of proteinase inhibitors in plant leaves. Proceedings of the National Academy of Sciences. 87: 7713–7716.

13. Fredericq, H. (1964) Influence formatrice de la lumière rouge-foncé sur le développement des thalles de *Marchantia polymorpha* L. [Formative influence of far-red light on the development of *Marchantia polymorpha* L. thalli]. Bull Soc Roy Bot Belgique. 98: 67–76.

14. Fredericq, H., and de Greef, J. (1966) Red (R), far-red (FR) photoreversible control of growth and chlorophyll content in light-grown thalli of *Marchantia polymorpha* L. Die Naturwissenschaften. 53: 337.

15. Harris, B.J., Clark, J.W., Schrempf, D., Szöllősi, G.J., Donoghue, P.C.J., Hetherington, A.M., et al. (2022) Divergent evolutionary trajectories of bryophytes and tracheophytes from a complex common ancestor of land plants. Nat Ecol Evol. 6: 1634–1643.

16. Hayashi, K., Horie, K., Hiwatashi, Y., Kawaide, H., Yamaguchi, S., Hanada, A., et al. (2010) Endogenous diterpenes derived from *ent*-kaurene, a common gibberellin precursor, regulate protonema differentiation of the moss *Physcomitrella patens*. Plant Physiology. 153: 1085–1097.

17. Hostettler, C., Kölling, K., Santelia, D., Streb, S., Kötting, O., and Zeeman, S.C. (2011) Analysis of Starch Metabolism in Chloroplasts. In Chloroplast Research in Arabidopsis: Methods and Protocols*, Volume II*. Edited by Jarvis, R.P. pp. 387–410 Humana Press, Totowa, NJ.

18. Inagaki, H., Miyamoto, K., Ando, N., Murakami, K., Sugisawa, K., Morita, S., et al. (2021) Deciphering OPDA Signaling Components in the Momilactone-Producing Moss *Calohypnum plumiforme*. Front Plant Sci. 12.

19. Inoue, K., Nishihama, R., Araki, T., and Kohchi, T. (2019) Reproductive induction is a far-red high irradiance response that is mediated by phytochrome and PHYTOCHROME INTERACTING FACTOR in *Marchantia polymorpha*. Plant and Cell Physiology. 60: 1136–1145.

20. Ishizaki, K., Chiyoda, S., Yamato, K.T., and Kohchi, T. (2008) *Agrobacterium*-mediated transformation of the haploid liverwort *Marchantia polymorpha* L., an emerging model for plant biology. Plant and Cell Physiology. 49: 1084–1091.

21. Ishizaki, K., Nishihama, R., Ueda, M., Inoue, K., Ishida, S., Nishimura, Y., et al. (2015) Development of Gateway binary vector series with four different selection markers for the liverwort *Marchantia polymorpha*. PLoS ONE. 10: e0138876.

22. Kato, M., and Mizuno, K. (2004) Caffeine synthase and related methyltransferases in plants. Front Biosci. 9: 1833–1842.

23. Kawamura, S., Romani, F., Yagura, M., Mochizuki, T., Sakamoto, M., Yamaoka, S., et al. (2022) MarpolBase Expression: a web-based, comprehensive platform for visualization and analysis of transcriptomes in the liverwort *Marchantia polymorpha*. Plant and Cell Physiology. 63: 1745–1755.

24. Koide, E., Suetsugu, N., Iwano, M., Gotoh, E., Nomura, Y., Stolze, S.C., et al. (2020) Regulation of photosynthetic carbohydrate metabolism by a Raf-like kinase in the liverwort *Marchantia polymorpha*. Plant and Cell Physiology. 61: 631–643.

25. Komatsu, A., Fujibayashi, M., Kumagai, K., Suzuki, H., Hata, Y., Takebayashi, Y., et al. (2025) KAI2-dependent signaling controls vegetative reproduction in Marchantia polymorpha through activation of LOG-mediated cytokinin synthesis. Nat Commun. 16: 1263.

26. Koo, Y.J., Kim, M.A., Kim, E.H., Song, J.T., Jung, C., Moon, J.-K., et al. (2007) Overexpression of salicylic acid carboxyl methyltransferase reduces salicylic acid-mediated pathogen resistance in *Arabidopsis thaliana*. Plant Mol Biol. 64: 1–15.

27. Kubota, A., Ishizaki, K., Hosaka, M., and Kohchi, T. (2013) Efficient *Agrobacterium*-mediated transformation of the liverwort *Marchantia polymorpha* using regenerating thalli. *Bioscience*, Biotechnology and Biochemistry. 77: 167–172.

28. Kumar, S., Kempinski, C., Zhuang, X., Norris, A., Mafu, S., Zi, J., et al. (2016) Molecular diversity of terpene synthases in the liverwort *Marchantia polymorpha*. The Plant Cell. 28: 2632–2650.

29. Lee, J.E., Goretti, D., Neumann, M., Schmid, M., and You, Y. (2020) A gibberellin methyltransferase modulates the timing of floral transition at the Arabidopsis shoot meristem. Physiol Plant. 170: 474–487.

30. Li, L., Zheng, Q., Jiang, W., Xiao, N., Zeng, F., Chen, G., et al. (2022) Molecular Regulation and Evolution of Cytokinin Signaling in Plant Abiotic Stresses. Plant Cell Physiol. 63: 1787–1805.

31. Love, M.I., Huber, W., and Anders, S. (2014) Moderated estimation of fold change and dispersion for RNA-seq data with DESeq2. Genome Biology. 15: 550.

32. Mashiguchi, K., Seto, Y., Onozuka, Y., Suzuki, S., Takemoto, K., Wang, Y., et al. (2022) A carlactonoic acid methyltransferase that contributes to the inhibition of shoot branching in Arabidopsis. Proceedings of the National Academy of Sciences. 119: e2111565119.

33. Miyazaki, S., Hara, M., Ito, S., Tanaka, K., Asami, T., Hayashi, K., et al. (2018) An ancestral gibberellin in a moss *Physcomitrella patens*. Molecular Plant. 11: 1097–1100.

34. Monte, I., Ishida, S., Zamarreño, A.M., Hamberg, M., Franco-Zorrilla, J.M., García-Casado, G., et al. (2018) Ligand-receptor co-evolution shaped the jasmonate pathway in land plants. Nat Chem Biol. 14: 480–488.

35. Morffy, N., and Strader, L.C. (2020) Old Town Roads: routes of auxin biosynthesis across kingdoms. Current Opinion in Plant Biology., Physiology and Metabolism 55: 21–27.

36. Morris, J.L., Puttick, M.N., Clark, J.W., Edwards, D., Kenrick, P., Pressel, S., et al. (2018) The timescale of early land plant evolution. Proceedings of the National Academy of Sciences. 115: E2274–E2283.

37. Naito, Y., Hino, K., Bono, H., and Ui-Tei, K. (2015) CRISPRdirect: software for designing CRISPR/Cas guide RNA with reduced off-target sites. Bioinformatics. 31: 1120–1123.

38. Patel, H., Manning, J., Ewels, P., Garcia, M.U., Peltzer, A., Hammarén, R., et al. (2025) nf-core/rnaseq: nf-core/rnaseq v3.22.1 - Polished Palladium Penguin (3.22.1). Zenodo. 10.5281/zenodo.17833254.

39. Patro, R., Duggal, G., Love, M.I., Irizarry, R.A., and Kingsford, C. (2017) Salmon provides fast and bias-aware quantification of transcript expression. Nature Methods. 14: 417–419.

40. Qin, G., Gu, H., Zhao, Y., Ma, Z., Shi, G., Yang, Y., et al. (2005) An Indole-3-Acetic Acid Carboxyl Methyltransferase Regulates Arabidopsis Leaf Development. Plant Cell. 17: 2693–2704.

41. Rademacher, W. (2000) Growth retardants : Effects on gibberellin biosynthesis and other metabolic pathways. Annual Review of Plant Physiology & Plant Molecular Biology. 51: 501–531.

42. Radhika, V., Ueda, N., Tsuboi, Y., Kojima, M., Kikuchi, J., Kudo, T., et al. (2015) Methylated Cytokinins from the Phytopathogen *Rhodococcus fascians* Mimic Plant Hormone Activity. Plant Physiol. 169: 1118–1126.

43. Ross, J.R., Nam, K.H., D’Auria, J.C., and Pichersky, E. (1999) *S*-Adenosyl-l-Methionine:Salicylic Acid Carboxyl Methyltransferase, an Enzyme Involved in Floral Scent Production and Plant Defense, Represents a New Class of Plant Methyltransferases. Archives of Biochemistry and Biophysics. 367: 9–16.

44. Seo, H.S., Song, J.T., Cheong, J.-J., Lee, Y.-H., Lee, Y.-W., Hwang, I., et al. (2001) Jasmonic acid carboxyl methyltransferase: A key enzyme for jasmonate-regulated plant responses. Proceedings of the National Academy of Sciences. 98: 4788–4793.

45. Shulaev, V., Silverman, P., and Raskin, I. (1997) Airborne signalling by methyl salicylate in plant pathogen resistance. Nature. 385: 718–721.

46. Soneson, C., Love, M.I., and Robinson, M.D. (2016) Differential analyses for RNA-seq: transcript-level estimates improve gene-level inferences. F1000Research. 1521.

47. Staswick, P.E., and Tiryaki, I. (2004) The Oxylipin Signal Jasmonic Acid Is Activated by an Enzyme That Conjugates It to Isoleucine in Arabidopsis[W]. Plant Cell. 16: 2117–2127.

48. Streubel, S., Deiber, S., Rötzer, J., Mosiolek, M., Jandrasits, K., and Dolan, L. (2023) Meristem dormancy in *Marchantia polymorpha* is regulated by a liverwort-specific miRNA and a clade III SPL gene. Current Biology. 33: 660–674.

49. Sugano, S.S., Shirakawa, M., Takagi, J., Matsuda, Y., Shimada, T., Hara-Nishimura, I., et al. (2014) CRISPR/Cas9-mediated targeted mutagenesis in the liverwort *Marchantia polymorpha* L. Plant and Cell Physiology. 55: 475–481.

50. Sun, R., Okabe, M., Miyazaki, S., Ishida, T., Mashiguchi, K., Inoue, K., et al. (2023) Biosynthesis of gibberellin-related compounds modulates far-red light responses in the liverwort *Marchantia polymorpha*. The Plant Cell. 35: 4111–4132.

51. Tanizawa, Y., Mochizuki, T., Yagura, M., Sakamoto, M., Fujisawa, T., Kawamura, S., et al. (2025) MarpolBase: Genome database for *Marchantia polymorpha* featuring high quality reference genome sequences. Plant Cell Physiol. pcaf159.

52. Tomizawa, Y., Minamino, N., Shimokawa, E., Kawamura, S., Komatsu, A., Hiwatashi, T., et al. (2023) Harnessing Deep Learning to Analyze Cryptic Morphological Variability of *Marchantia polymorpha*. Plant Cell Physiol. 64: 1343–1355.

53. Varbanova, M., Yamaguchi, S., Yang, Y., McKelvey, K., Hanada, A., Borochov, R., et al. (2007) Methylation of gibberellins by *Arabidopsis* GAMT1 and GAMT2. The Plant Cell. 19: 32–45.

54. Wakabayashi, T., Yasuhara, R., Miura, K., Takikawa, H., Mizutani, M., and Sugimoto, Y. (2021) Specific methylation of (11R)-carlactonoic acid by an Arabidopsis SABATH methyltransferase. Planta. 254: 88.

55. Wang, W., Guo, H., Bowman, J.L., and Chen, F. (2024) Plant SABATH Methyltransferases: Diverse Functions, Unusual Reaction Mechanisms and Complex Evolution. Critical Reviews in Plant Sciences. 43: 291–312.

56. Ward, L.C., McCue, H.V., and Carnell, A.J. (2021) Carboxyl Methyltransferases: Natural Functions and Potential Applications in Industrial Biotechnology. ChemCatChem. 13: 121–128.

57. Xing, S., Qin, G., Shi, Y., Ma, Z., Chen, Z., Gu, H., et al. (2007) GAMT2 Encodes a Methyltransferase of Gibberellic Acid That is Involved in Seed Maturation and Germination in Arabidopsis. Journal of Integrative Plant Biology. 49: 368–381.

58. Yu, G., Wang, L.-G., Han, Y., and He, Q.-Y. (2012) clusterProfiler: an R Package for Comparing Biological Themes Among Gene Clusters. OMICS: A Journal of Integrative Biology. 16: 284–287.

59. Zhang, C., Chaiprasongsuk, M., Chanderbali, A.S., Chen, X., Fu, J., Soltis, D.E., et al. (2020) Origin and evolution of a gibberellin-deactivating enzyme GAMT. Plant Direct. 4: e00287.

60. Zhang, C., Chen, X., Crandall-Stotler, B., Qian, P., Köllner, T.G., Guo, H., et al. (2019) Biosynthesis of methyl (E)-cinnamate in the liverwort *Conocephalum salebrosum* and evolution of cinnamic acid methyltransferase. Phytochemistry. 164: 50–59.

61. Zhao, N., Boyle, B., Duval, I., Ferrer, J.-L., Lin, H., Seguin, A., et al. (2009) SABATH methyltransferases from white spruce (*Picea glauca*): gene cloning, functional characterization and structural analysis. Tree Physiol. 29: 947–957.

62. Zhao, N., Ferrer, J.-L., Moon, H.S., Kapteyn, J., Zhuang, X., Hasebe, M., et al. (2012) A SABATH Methyltransferase from the moss *Physcomitrella patens* catalyzes *S*-methylation of thiols and has a role in detoxification. Phytochemistry. 81: 31–41.

63. Zubieta, C., Ross, J.R., Koscheski, P., Yang, Y., Pichersky, E., and Noel, J.P. (2003) Structural Basis for Substrate Recognition in the Salicylic Acid Carboxyl Methyltransferase Family. Plant Cell. 15: 1704–1716.

