## Supplemental Data for "A SABATH family enzyme regulates development via the gibberellin-related pathway in the liverwort *Marchantia polymorpha*"

**Supplementary Data S1.**

Unrooted phylogenetic tree of SABATH family genes, in the full form. Numbers next to internal nodes represent percentage support values from 1000 bootstraps by RAxML-NG.

- Streptophyte Algae
- Liverwort
  - *M. polymorpha*
- Hornwort
- Moss
- Lycophyte
- Fern
- Gymnosperm
- Basal Angiosperm
- Monocot
- Eudicot

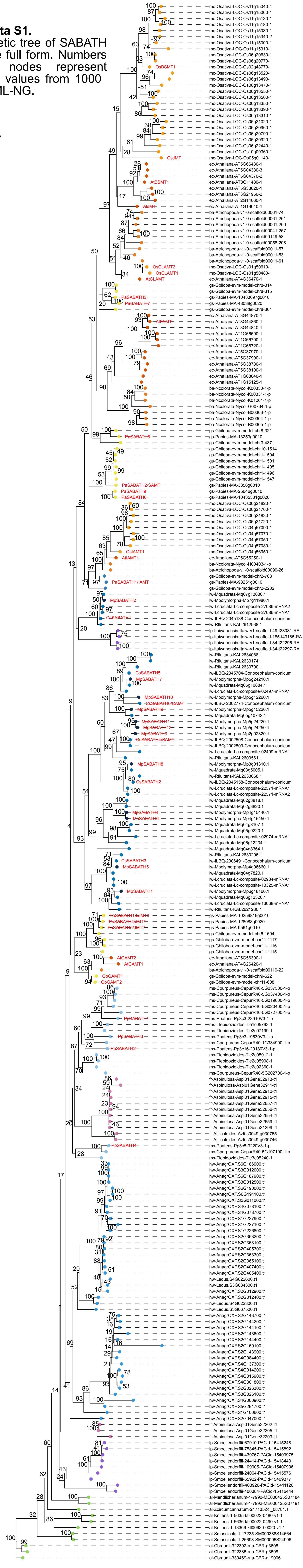
