## Supplemental FiguresTables for "A SABATH family enzyme regulates development via the gibberellin-related pathway in the liverwort *Marchantia polymorpha*"

**Supplementary Figures S1-S9**

**Supplementary Tables S1-S3**

**Supplementary References**

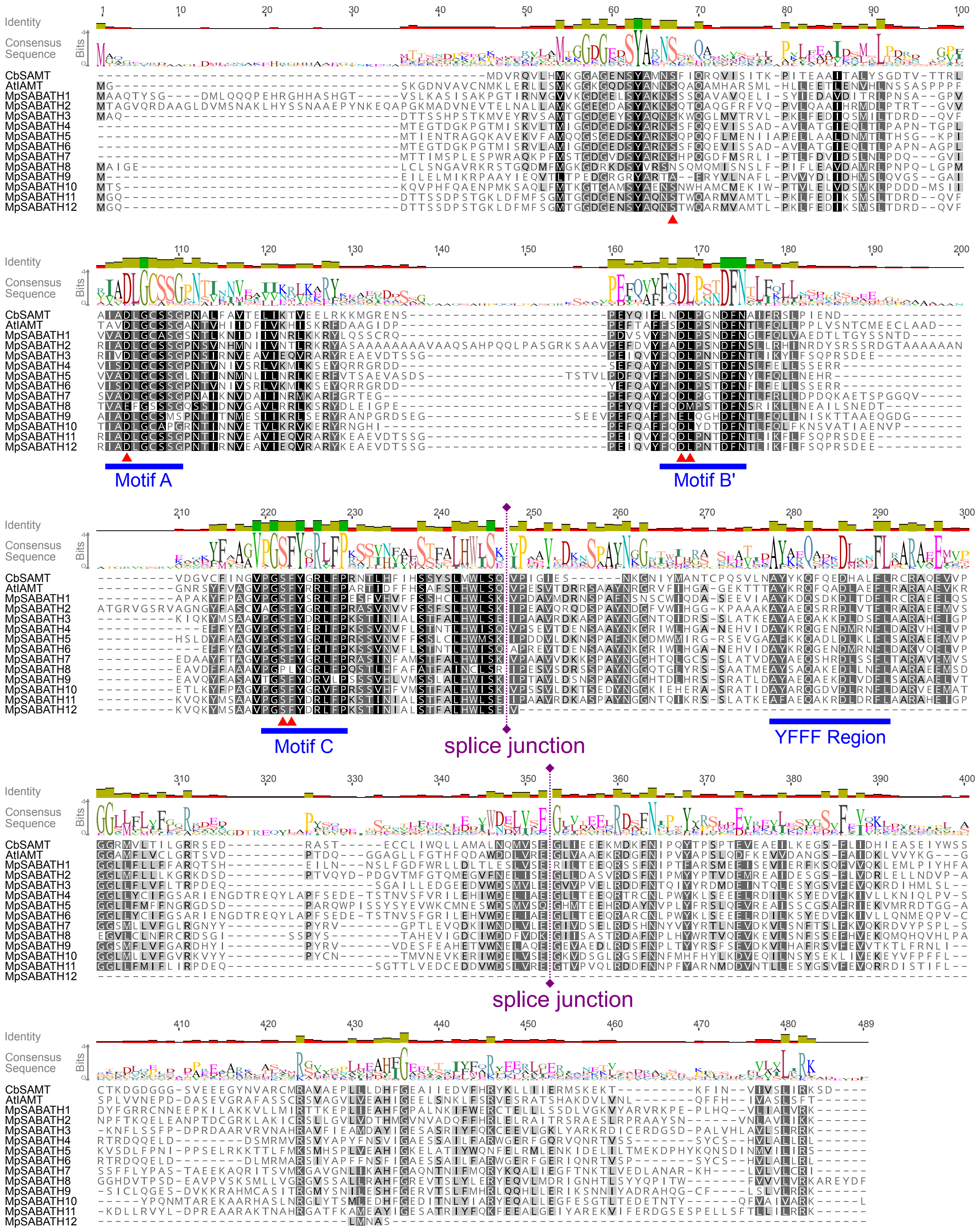


**Supplementary Figure S1.** Multiple sequence alignments of SABATH family proteins in *M. polymorpha*

The sequences of SABATH family proteins in *M. polymorpha* are aligned with two proteins with determined protein structure, *Clarkia breweri* salicylic acid methyltransferase (CbSAMT) (Zubieta et al.,

2003) and *Arabidopsis thaliana* IAA methyltransferase 1 (AtIAMT1) (Zhao et al., 2008) using MAFFT. The residues are shaded according to similarity, and the consensus sequence is depicted with sequence logo. Motif A, Motif B', Motif C, and YFFF regions are indicated with blue lines. SAM-binding residues inferred from the structure of CbSAMT (Zubieta et al., 2003) are indicated by red triangles. The two conserved splice junction sites are indicated by purple dashed lines.

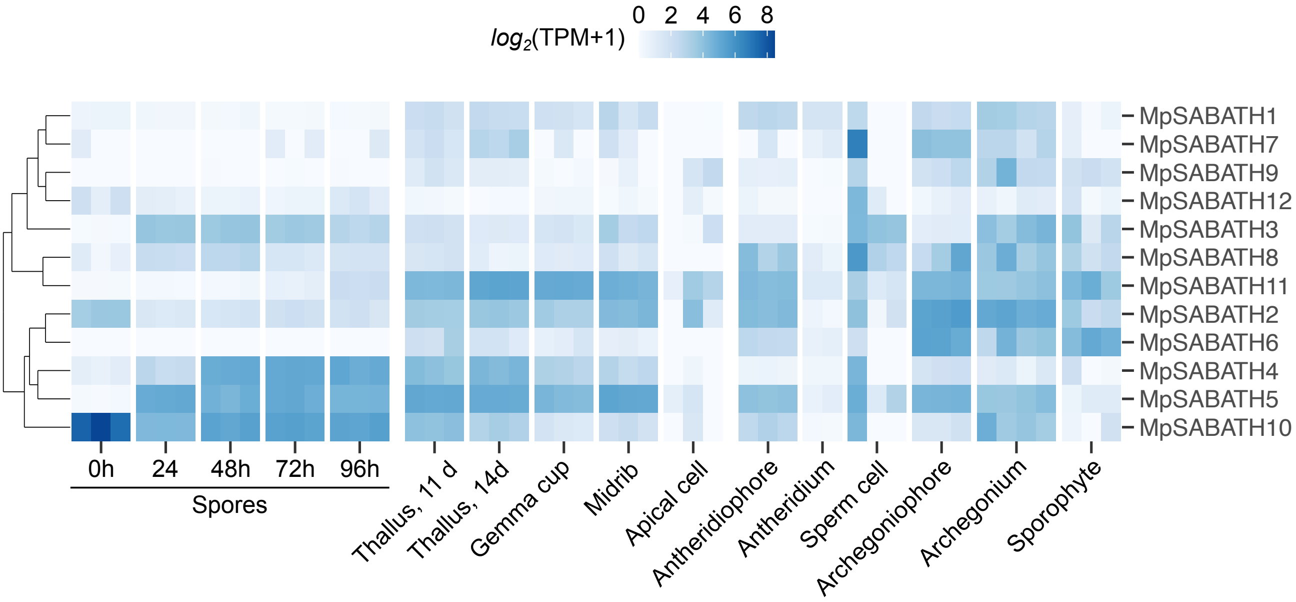


**Supplementary Figure S2.** Expression pattern of Mp*SABATH* genes, based on transcriptome data retrieved from MarpolBase Expression (https://mbex.marchantia.info/) (Kawamura et al., 2022).

The original data sources are: spores collected 0-72 hrs post germination (Bowman et al., 2017), 11-day-old whole thallus (Higo et al., 2016), 14-day-old thallus (Karaaslan et al., 2020), gemma cup and mid rib regions from 21-d-old thallus (Yasui et al., 2019), cells around the apical meristem (Frank and Scanlon, 2015), antheridiophore receptacles and antheridia (Higo et al., 2016), sperm cells (Julca et al., 2021), archegoniophore receptacles (Higo et al., 2016), archegonia (Hisanaga et al., 2021), sporophytes (Frank and Scanlon, 2015).

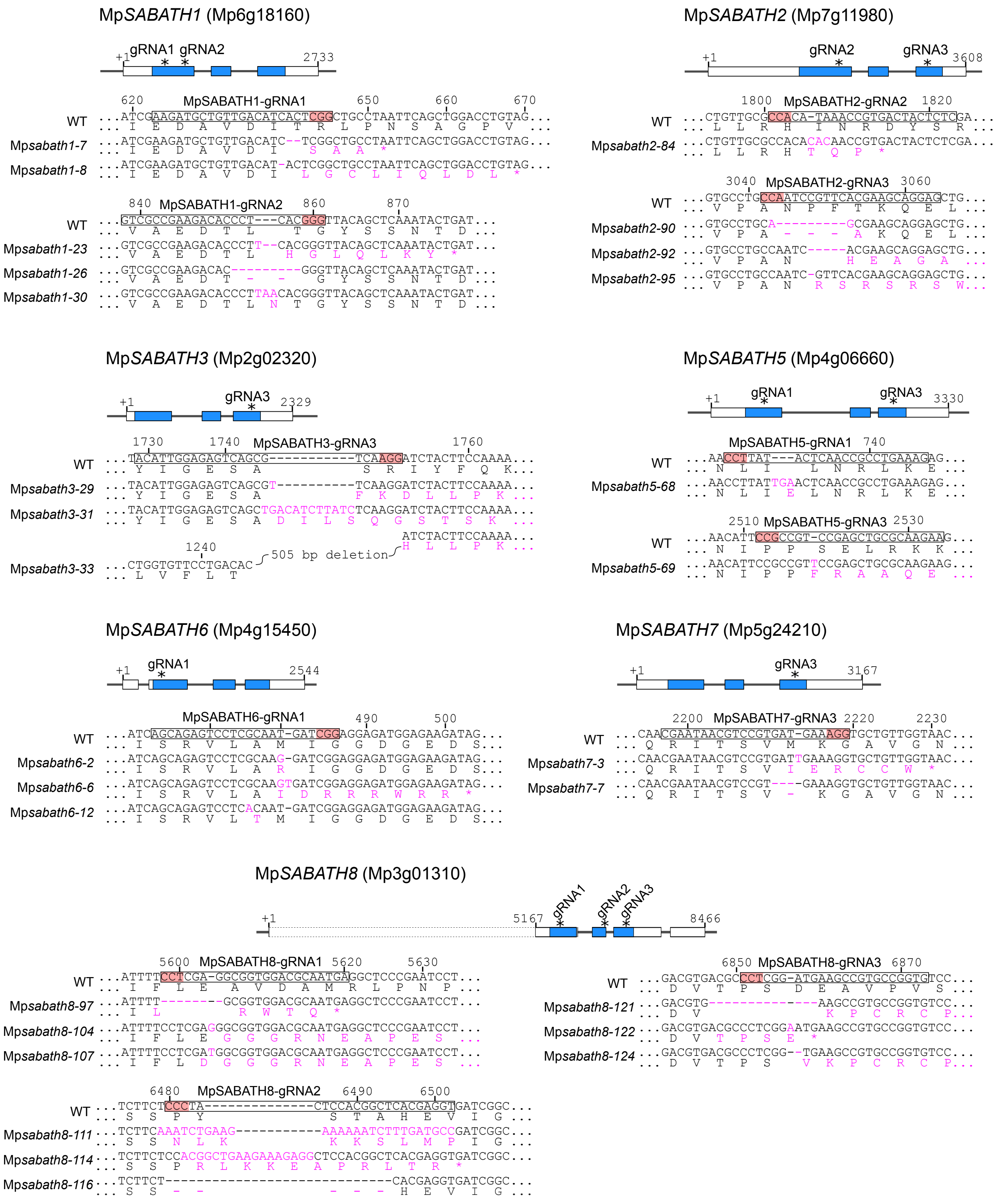


(To be continued)

(Continued)

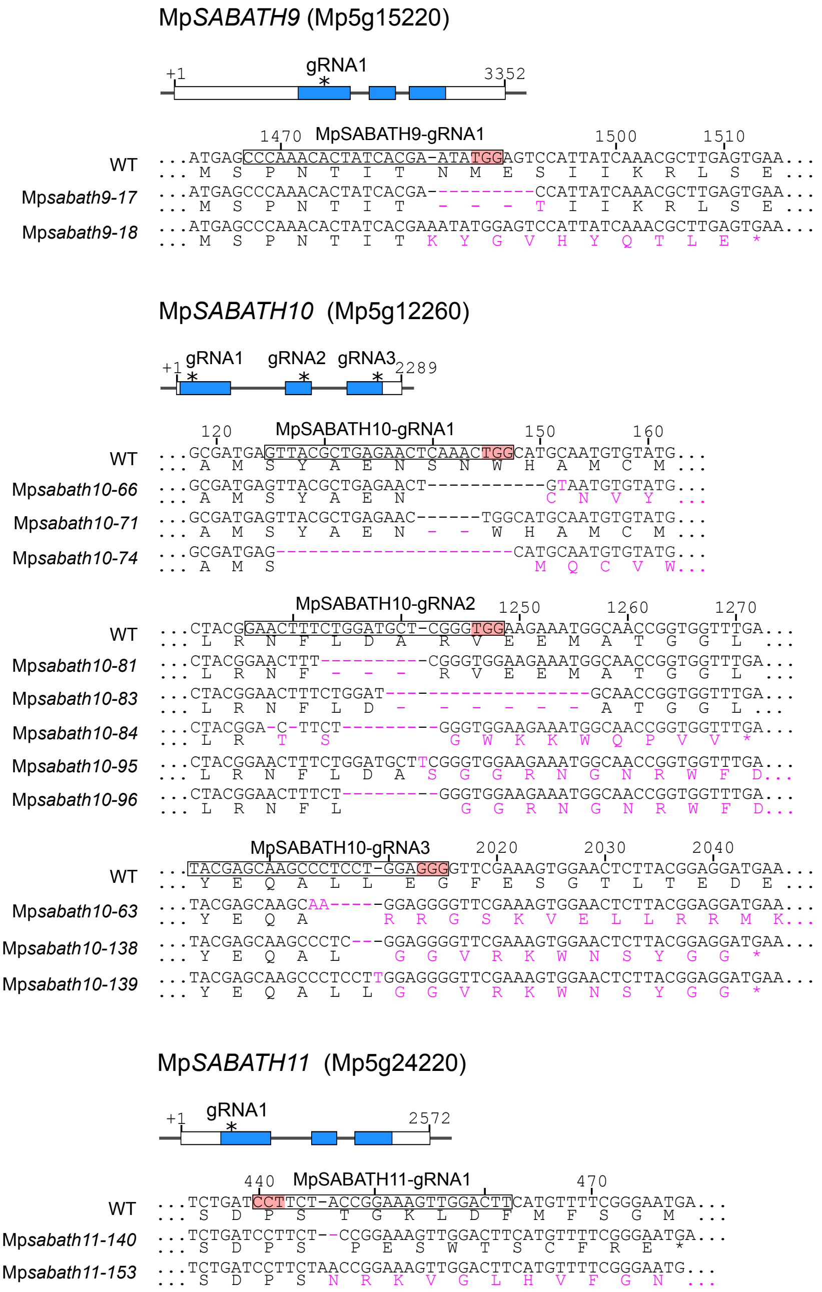


**Supplemental Figure S3.** Genotypes of Mp*sabath* mutants obtained in the initial screening.

For Mp*SABATH1*, Mp*SABATH2*, Mp*SABATH5*, Mp*SABATH6*, Mp*SABATH8*, Mp*SABATH9*, Mp*SABATH10* and Mp*SABATH11*, at least one frame-shift mutation was introduced near the N-terminus, likely causing loss of gene function. For MpSABATH3, although all the mutations were obtained with gRNA3 close to the C-terminus, Mp*sabath3-33* carries a 505-bp deletion which is expected to severely affect the gene function. For Mp*SABATH7*, two mutants were obtained with gRNA3, causing frameshift or 1-aa deletion near the C-terminus. No mutant was obtained for *MpSABATH4*.

For Mp*SABATH8*, we manually curated the transcription start site (TSS) was manually curated based on the transcriptome coverage and cap analysis of gene expression (CAGE) data on https://marchantia.info/, as shown in Figure1. However, for the convenience to compare with the current standard reference genome, here we labeled the TSS from MpTak_ver7.1 was labeled as +1 in this figure.

In the schematic presentations of genomic structures, white and blue rectangles represent untranslated and coding regions of exons, respectively. Targets of guide RNAs are indicated by asterisks (*) in the scheme, and marked with frames in the sequences (pink shading: the protospacer adjacent motif). Indels and substitutions are shown in magenta letters. Numbers above the sequences indicate positions relative to the TSS (+1).

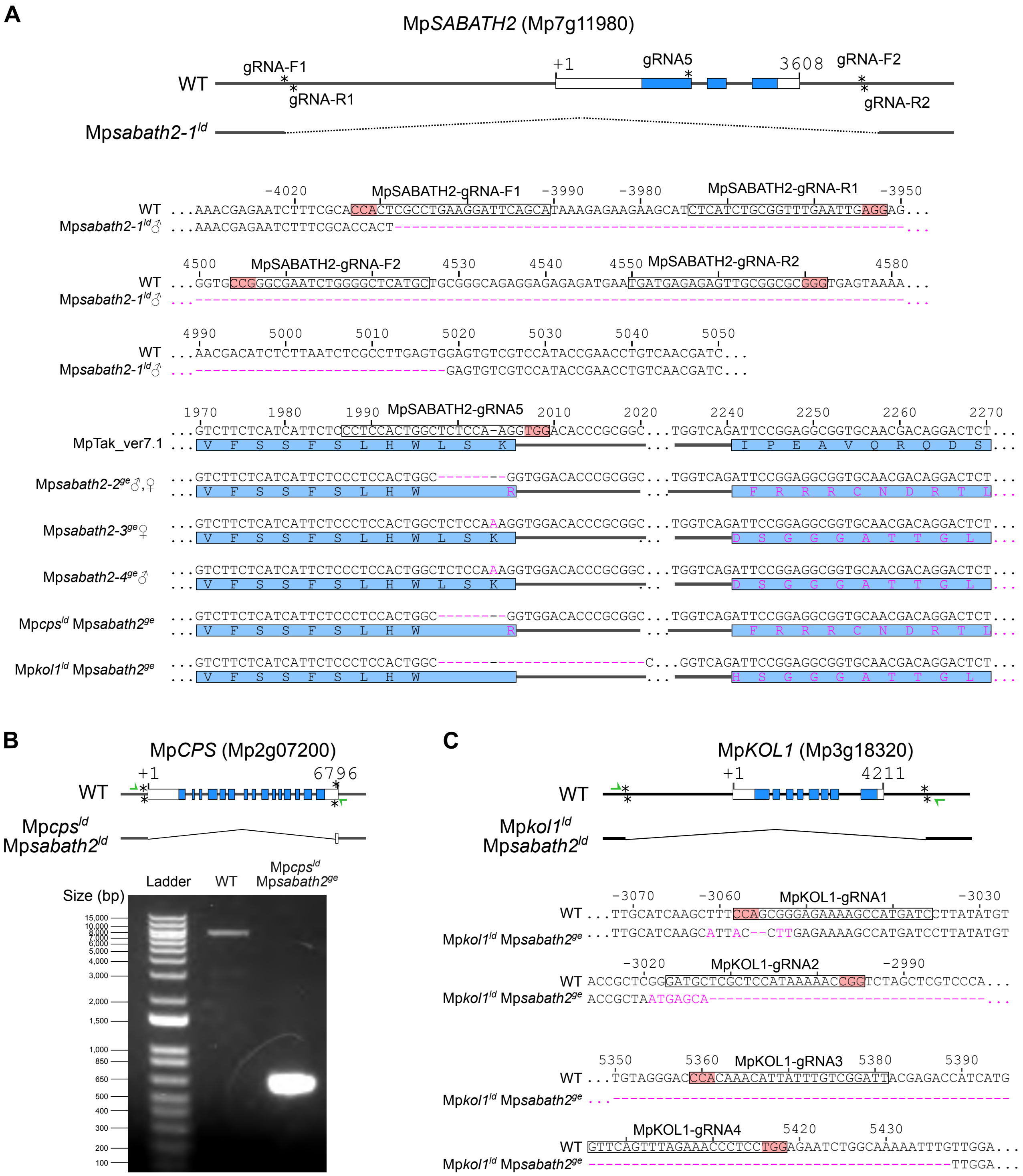


**Supplemental Figure S4.** Genotype of Mp*sabath2* mutants created after the initial screening.

A, Genotype information for mutations in the Mp*SABATH2* locus. Mp*sabath2-2*$^{ge}$♂, Mp*sabath2-3*$^{ge}$♀ and Mp*sabath2-4*$^{ge}$♂ are independent alleles obtained from different transformants, although Mp*sabath2-3*$^{ge}$♀ and Mp*sabath2-4*$^{ge}$♂ happen to carry the same type of mutation by coincidence. Mp*sabath2-2*$^{ge}$♀ is the female seggregant obtained by backcrossing Mp*sabath2-2*$^{ge}$♂ to the WT, and

Mp*cps*$^{ld}$ Mp*sabath2*$^{ge}$ was created from Mp*sabath2-2*$^{ge}$♂ by thallus transformation. Mp*kol1*$^{ld}$ Mp*sabath2*$^{ge}$ was created from Mp*kol1*$^{ld}$ so it carries another type of Mp*sabath2* mutation.

B, Genotyping PCR with the primers MpCPS-gt-F and MpCPS-gt-R, showing the deletion of the Mp*CPS* coding region confirming the deletion in Mp*cps*$^{ld}$ Mp*sabath2*$^{ge}$.

C, Genotype information of the Mp*kol1* mutation in Sanger-sequence result of Mp*KOL1* region confirming the deletion in Mp*kol1*$^{ld}$ Mp*sabath2*$^{ge}$. We followed the method described in our previous publication for panel B and C (Sun et al. 2023).

In the schematic presentations of genomic structures, white and blue rectangles represent untranslated and coding regions of exons, respectively. Targets of guide RNAs are indicated by asterisk (*) in the scheme, and marked with frames in the sequence (pink shading: the protospacer adjacent motif). Indels and substitutions are shown in magenta letters. Numbers above the sequences indicate positions relative to the TSS (+1). In (B) and (C), green arrows indicate the binding sites of genotyping primers.

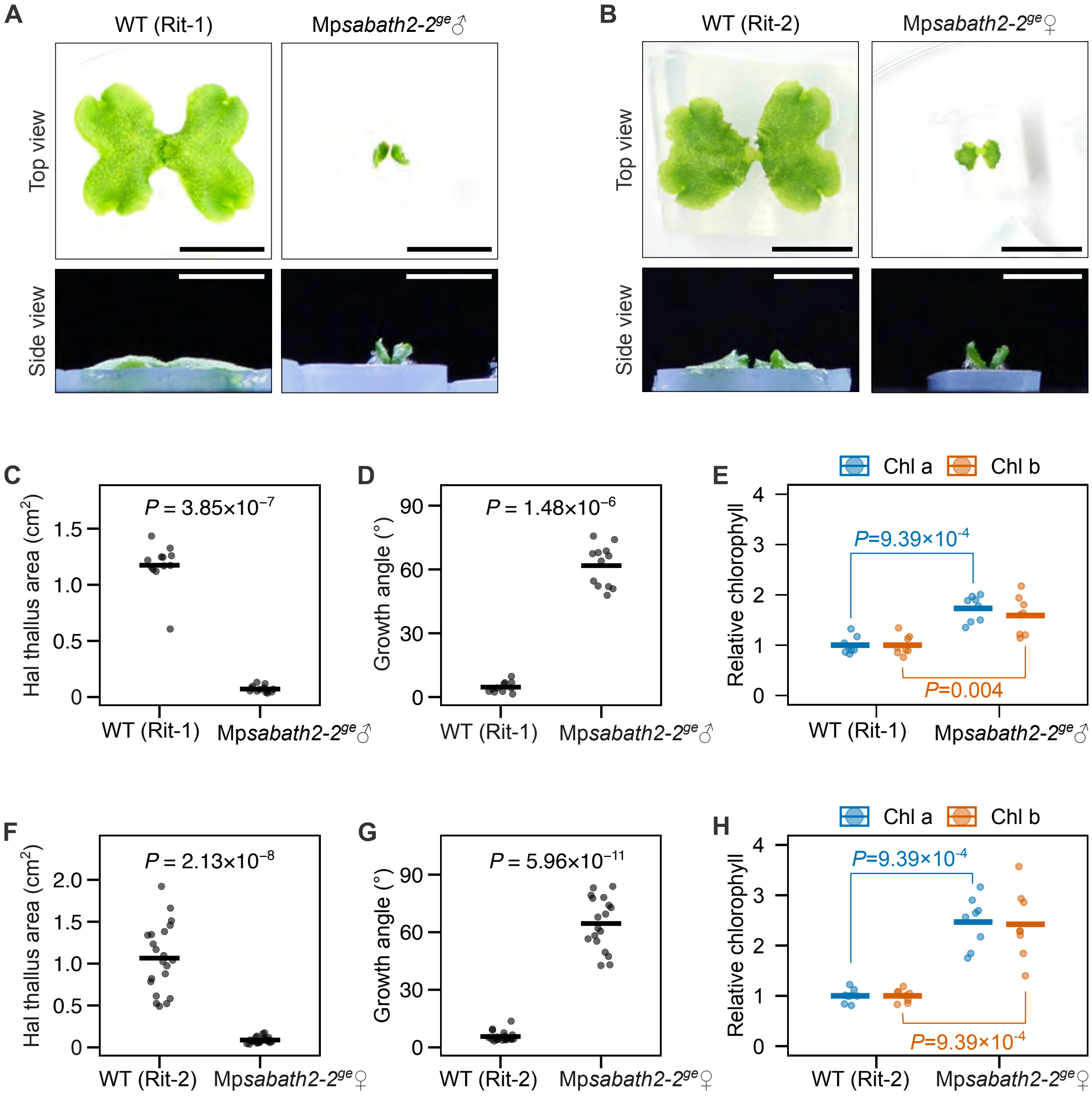


**Supplemental Figure S5.** Phenotypes of male and female Mp*sabath2-2*$^{ge}$ plants under cW condition.

A-B, Photos of 14-day-old Mp*sabath2-2*$^{ge}$ plants grown from gemmae under cW conditions. Bars = 1 cm.

C-H, Measurements of the thallus size (C: male, F: female), growth angle (D: male, G: female), and relative chlorophyll level (E: male, H: female) in 14-day-old plants. The *P* values were calculated by the Mann-Whitney *U* test (n=11-12 for C-D, n= 21-22 for F-G, n=8 for E and H).

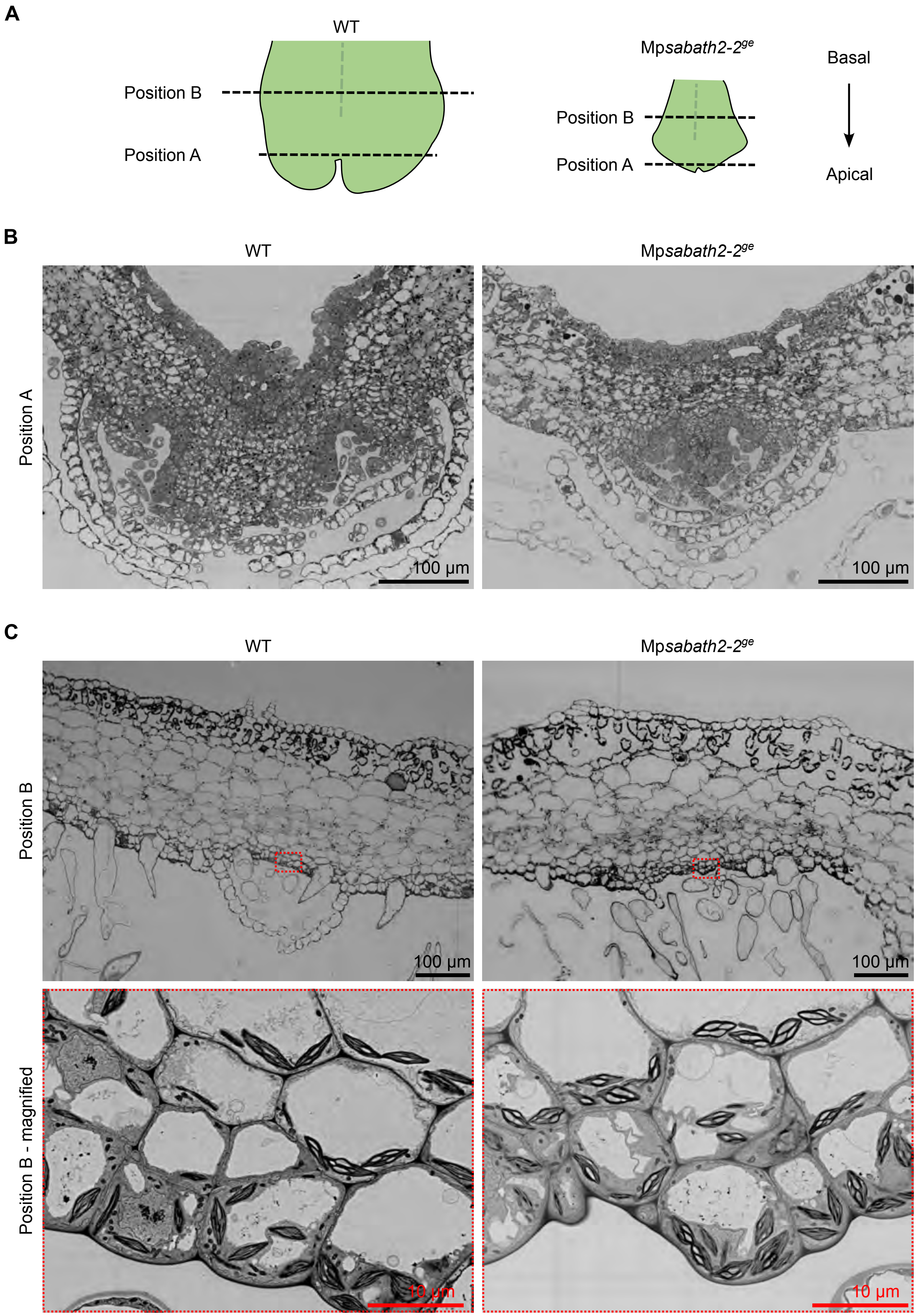
A
WT
Position B
Position A
Mpsabath2-2ge
Position B
Position A
Basal
Apical
B
WT
Mpsabath2-2ge
Position A
100 µm
100 µm
C
WT
Mpsabath2-2ge
Position B
100 µm
100 µm
Position B - magnified
10 µm
10 µm

**Supplemental Figure S6**. Ultra-thin cross sections of WT and Mp*sabath2-2[ge]* plants.

A, Illustration showing the approximate positions of sections in 14-day-old plants grown from gemmae under cW conditions.

B, Sections near the apical region. No obvious cell size reduction was observed in Mp*sabath2-2[ge]* plants; instead, the number of cells seemed to be decreased in the mutant. Bars = 100 μm.

C, Sections near the middle of the thallus. Mp*sabath2-2[ge]* plants showed increased starch accumulation in the chloroplasts from the ventral cell layers near the midrib. Bars = 100 μm for top images, 10 μm for bottom images.

Images in (B-C) were taken with field emission scanning electron microscopes (FE-SEM).

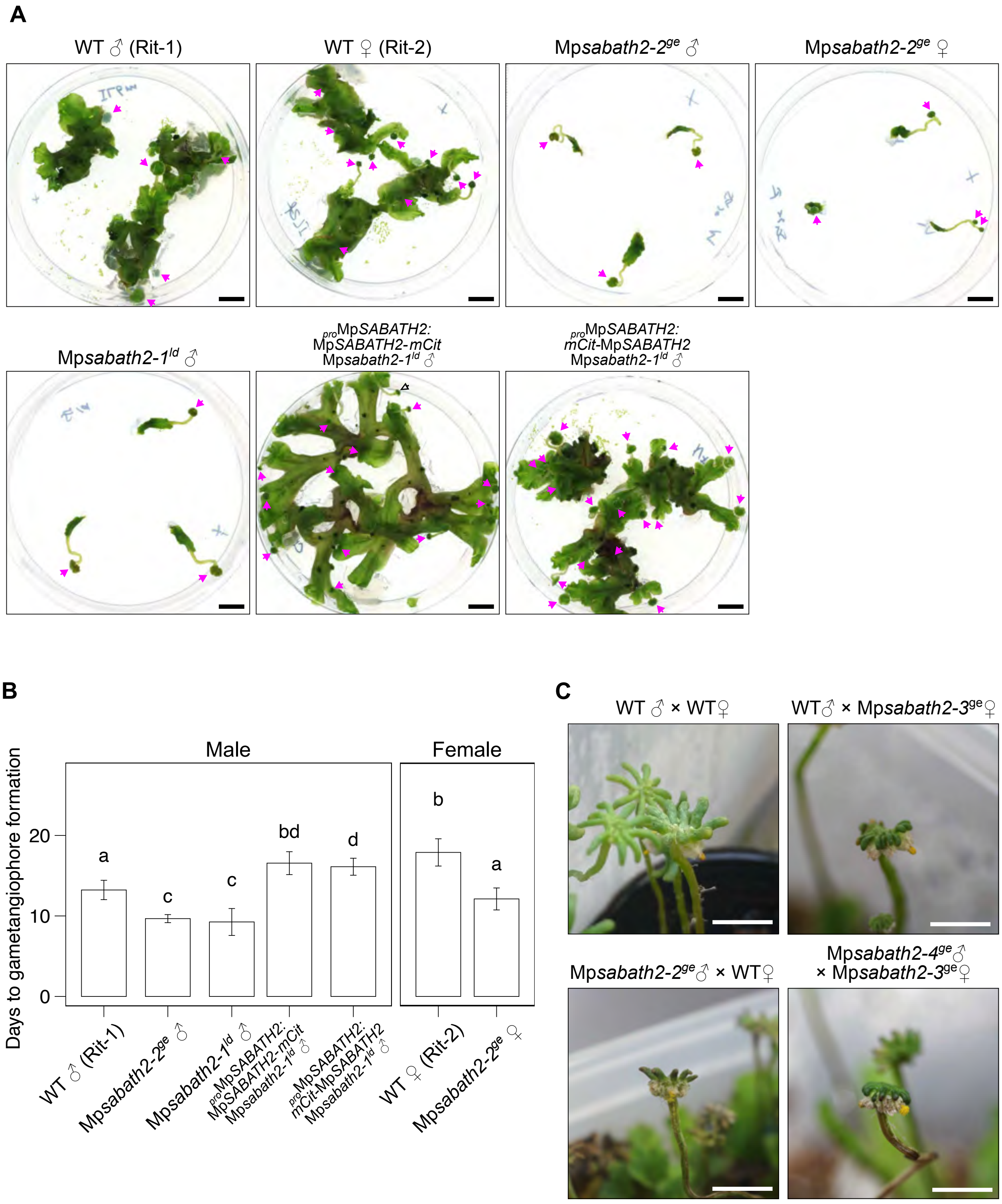


**Supplemental Figure S7**. Sexual reproduction of Mp*sabath2* mutants under FR-enriched inductive conditions (cW+FR).

A, Photos of 31-day-old plants. The plants were grown from gemmae under cW conditions for 10 days, then half-thallus fragments were cultured under cW+FR conditions for 21 days. Magenta arrows indicate the receptacles of gametangiophores. Bars = 1 cm.

B, Days to first gametangiophore formation after FR irradiation. Non-overlapping letters indicate significant statistical difference by pairwise Mann-Whitney *U* test with Benjamini-Hochberg adjustment (n=8-9).

C, Photos of female receptacles, showing the formation of mature sporangia from crossing experiments among WT and Mp*sabath2*$^{ge}$ plants. Bars = 1 cm.

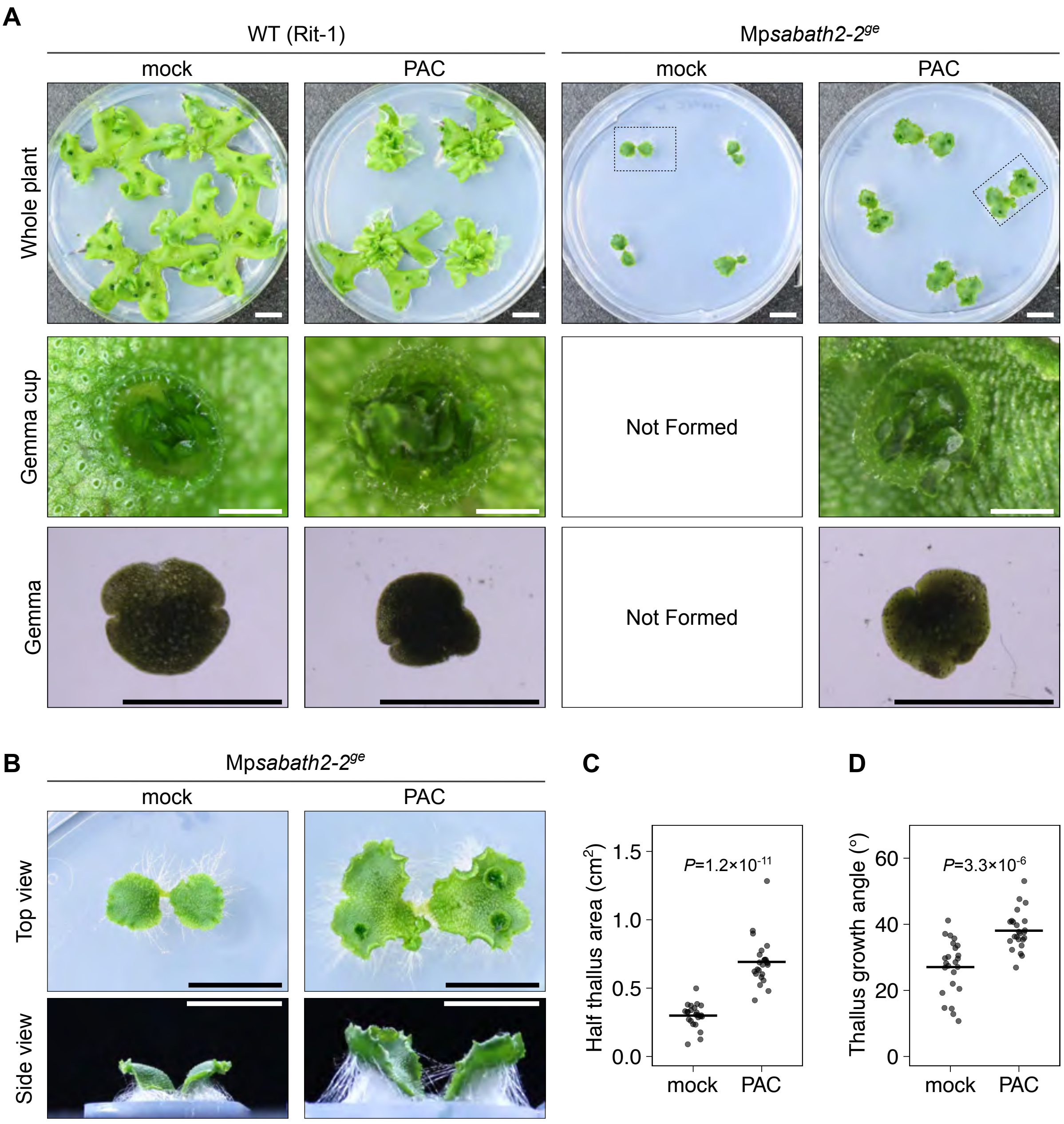


**Supplemental Figure S8.** Inhibition of diterpenoid biosynthesis pathway by PAC treatment suppressed Mp*sabath2* phenotypes.

A, Phenotypes of 21-day-old plants grown from gemmae under cW conditions with or without 20 μM PAC, showing the morphology of thallus, gemma cup, and gemma. Bars= 1 cm for thallus, 1 mm for gemma cups and gemmae.

B, Close-view of of Mp*sabath2*$^{ge}$ plants indicated by dashed frame in (A), showing the formation of gemma cups under PAC treatment.

C-D, Measurements of thallus area (C) and growth angle (B) for Mp*sabath2-2*$^{ge}$ plants shown in (A) and (B). The *P* values was calculated by Welch's *t*-test (n=24).

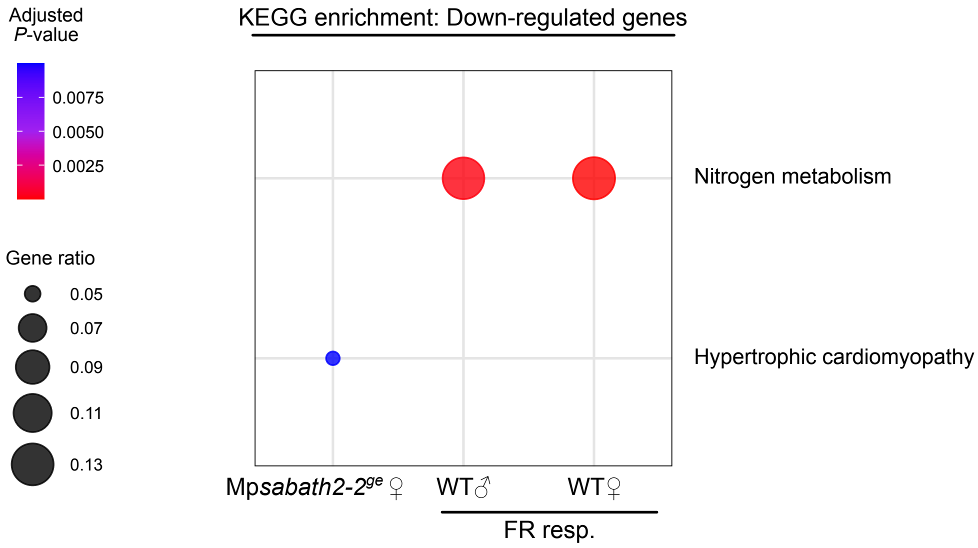


**Supplemental Figure S9**. KEGG enrichment analysis of down-regulated genes

No common enriched terms were found between Mp*sabath2*$^{ge}$ and FR response in WT plants.

### Supplemental Tables

**Supplemental Table S1.** List of major genes involved in this research

| Gene Name | MpGeneID | JGI3.1 GeneID |
|---|---|---|
| MpSABATH1 | Mp6g18160 | Mapoly0038s0025 |
| MpSABATH2 | Mp7g11980 | Mapoly0003s0211 |
| MpSABATH3 | Mp2g02320 | Mapoly0130s0039 |
| MpSABATH4 | Mp4g15440 | Mapoly0054s0007 |
| MpSABATH5 | Mp4g06660 | Mapoly0125s0011 |
| MpSABATH6 | Mp4g15450 | Mapoly0054s0010 |
| MpSABATH7 | Mp5g24210 | Mapoly0010s0035 |
| MpSABATH8 | Mp3g01310 | Mapoly0007s0125 |
| MpSABATH9 | Mp5g15220 | Mapoly0071s0088 |
| MpSABATH10 | Mp5g12260 | Mapoly0092s0080 |
| MpSABATH11 | Mp5g24220 | Mapoly0010s0033 |
| MpSABATH12 | Mp5g24250 | - |
| MpCPS | Mp2g07200 | Mapoly0015s0008 |
| MpKOL1 | Mp3g18320 | Mapoly0140s0010 |
| MpKS | Mp6g05950 | Mapoly0097s0049 |
| MpTPS1 | Mp6g05430 | Mapoly0167s0025 |
| MpTPS5 | Mp7g00020 | Mapoly0046s0122 |
| MpTPS6 | Mp3g13150 | Mapoly0050s0107 |
| MpKAOL1 | Mp4g23680 | Mapoly0020s0131 |
| MpKAOL2 | Mp1g25410 | Mapoly0002s0331 |
| MpKAOL3 | Mp2g10420 | Mapoly0023s0011 |
| MpDOXC14 | Mp3g01290 | Mapoly0007s0123 |
| MpDOXC22 | Mp3g25160 | Mapoly0100s0029 |

**Supplemental Table S2**. List of primers used in this study

| Name | Oligo Sequence (5′ -> 3′) | Used for |
|---|---|---|
| Mpsabath1-gRNA1-Fw | ctcgAAGATGCTGTTGACATCACT | Genome editing of Mp*SABATH1* |
| Mpsabath1-gRNA1-Rv | aaacAGTGATGTCAACAGCATCTT | |
| Mpsabath1-gRNA2-Fw | ctcgGTCGCCGAAGACACCCTCAC | |
| Mpsabath1-gRNA2-Rv | aaacGTGAGGGTGTCTTCGGCGAC | |
| Mpsabath1-gRNA3-Fw | ctcgATTTCGCCGACTACTTCGGA | |
| Mpsabath1-gRNA3-Rv | aaacTCCGAAGTAGTCGGCGAAAT | |
| Mpsabath2-gRNA1-Fw | ctcgGGCGCTTGCTCCTTGTTGTA | Genome editing of Mp*SABATH2* (intial screening) |
| Mpsabath2-gRNA1-Rv | aaacTACAACAAGGAGCAAGCGCC | |
| Mpsabath2-gRNA2-Fw | ctcgGAGAGTAGTCACGGTTTATG | |
| Mpsabath2-gRNA2-Rv | aaacCATAAACCGTGACTACTCTC | |
| Mpsabath2-gRNA3-Fw | ctcgCTCCTGCTTCGTGAACGGAT | |
| Mpsabath2-gRNA3-Rv | aaacATCCGTTCACGAAGCAGGAG | |
| MpSABATH2-gRNA_F1_A | ctcgTGCTGAATCCTTCAGGCGAG | Genome editing of Mp*SABATH2* (large deletion) |
| MpSABATH2-gRNA_F1_B | aaacCTCGCCTGAAGGATTCAGCA | |
| MpSABATH2-gRNA_F2_A | ctcgGCATGAGCCCCAGATTCGCC | |
| MpSABATH2-gRNA_F2_B | aaacGGCGAATCTGGGGCTCATGC | |
| MpSABATH2-gRNA_R1_A | ctcgCTCATCTGCGGTTTGAATTG | |
| MpSABATH2-gRNA_R1_B | aaacCAATTCAAACCGCAGATGAG | |
| MpSABATH2-gRNA_R2_A | ctcgTGATGAGAGAGTTGCGGCGC | |
| MpSABATH2-gRNA_R2_B | aaacGCGCCGCAACTCTCTCATCA | |
| MpSABATH2_N1 | CTCGTCAGTTTTGTAAGGGC | Genotyping of Mp*sabath2ld* |
| MpSABATH2_N_sequence | GGTTGCGAACATGGGAGTA | |
| MpSABATH2-gRNA5_nb | ctcgCCTCCACTGGCTCTCCAAGG | Genome editing of Mp*SABATH2* (others) |
| MpSABATH2-gRNA5_nf | aaacCCTTGGAGAGCCAGTGGAGG | |
| MpSABATH2-gRNA5_primer | CCGAAAATCTGCGGCTGTAC | Genotyping of Mp*sabath2*$^{ge}$ created with gRNA5 |
| MpSABATH2-gRNA5_primer2 | AAGCCATCGTTGTAAGCTGG | |
| Mpsabath3-gRNA1-Fw | ctcgAGAGATACAAGTCTACTTCC | Genome editing of Mp*SABATH3* |
| Mpsabath3-gRNA1-Rv | aaacGGAAGTAGACTTGTATCTCT | |
| Mpsabath3-gRNA2-Fw | ctcgTCAAATAGATCGATCAAGCT | |
| Mpsabath3-gRNA2-Rv | aaacAGCTTGATCGATCTATTTGA | |
| Mpsabath3-gRNA3-Fw | ctcgACATTGGAGAGTCAGCGTCA | |
| Mpsabath3-gRNA3-Rv | aaacTGACGCTGACTCTCCAATGT | |
| Mpsabath4-gRNA1-Fw | ctcgACTCTGCCTGCTCCAAACAC | Genome editing of Mp*SABATH4* |
| Mpsabath4-gRNA1-Rv | aaacGTGTTTGGAGCAGGCAGAGT | |
| Mpsabath4-gRNA2-Fw | ctcgGGCAGGGAGAAAATGACATG | |
| Mpsabath4-gRNA2-Rv | aaacCATGTCATTTTCTCCCTGCC | |
| Mpsabath4-gRNA3-Fw | ctcgGGGGAGAACGATTCGGACAA | |
| Mpsabath4-gRNA3-Rv | aaacTTGTCCGAATCGTTCTCCCC | |
| Mpsabath5-gRNA1-Fw | ctcgCTTTCAGGCGGTTGAGTATA | Genome editing of Mp*SABATH5* |
| Mpsabath5-gRNA1-Rv | aaacTATACTCAACCGCCTGAAAG | |

| Name | Oligo Sequence (5′ -> 3′) | Used for |
|---|---|---|
| Mpsabath5-gRNA2-Fw | ctcgGTCGCCCTTATTGAAAGCCG | Genome editing of Mp*SABATH5* |
| Mpsabath5-gRNA2-Rv | aaacCGGCTTTCAATAAGGGCGAC | |
| Mpsabath5-gRNA3-Fw | ctcgTTCTTGCGCAGCTCGGACGG | |
| Mpsabath5-gRNA3-Rv | aaacCCGTCCGAGCTGCGCAAGAA | |
| Mpsabath6-gRNA1-Fw | ctcgAGCAGAGTCCTCGCAATGAT | Genome editing of Mp*SABATH6* |
| Mpsabath6-gRNA1-Rv | aaacATCATTGCGAGGACTCTGCT | |
| Mpsabath6-gRNA2-Fw | ctcgACACATTCACTGACGACTTG | |
| Mpsabath6-gRNA2-Rv | aaacCAAGTCGTCAGTGAATGTGT | |
| Mpsabath6-gRNA3-Fw | ctcgTTCTCATCTGTGACTTCGCG | |
| Mpsabath6-gRNA3-Rv | aaacCGCGAAGTCACAGATGAGAA | |
| Mpsabath7-gRNA1-Fw | ctcgATGTCGACGCTATCATTAAC | Genome editing of Mp*SABATH7* |
| Mpsabath7-gRNA1-Rv | aaacGTTAATGATAGCGTCGACAT | |
| Mpsabath7-gRNA2-Fw | ctcgAGAGCGGTCGAAATGGTTTC | |
| Mpsabath7-gRNA2-Rv | aaacGAAACCATTTCGACCGCTCT | |
| Mpsabath7-gRNA3-Fw | ctcgCGAATAACGTCCGTGATGAA | |
| Mpsabath7-gRNA3-Rv | aaacTTCATCACGGACGTTATTCG | |
| Mpsabath8-gRNA1-Fw | ctcgTCATTGCGTCCACCGCCTCG | Genome editing of Mp*SABATH8* |
| Mpsabath8-gRNA1-Rv | aaacCGAGGCGGTGGACGCAATGA | |
| Mpsabath8-gRNA2-Fw | ctcgACCTCGTGAGCCGTGGAGTA | |
| Mpsabath8-gRNA2-Rv | aaacTACTCCACGGCTCACGAGGT | |
| Mpsabath8-gRNA3-Fw | ctcgCACCGGCACGGCTTCATCCG | |
| Mpsabath8-gRNA3-Rv | aaacCGGATGAAGCCGTGCCGGTG | |
| Mpsabath9-gRNA1-Fw | ctcgCCCAAACACTATCACGAATA | Genome editing of Mp*SABATH9* |
| Mpsabath9-gRNA1-Rv | aaacTATTCGTGATAGTGTTTGGG | |
| Mpsabath9-gRNA2-Fw | ctcgCAATATCTCTCTTACCTGGA | |
| Mpsabath9-gRNA2-Rv | aaacTCCAGGTAAGAGAGATATTG | |
| Mpsabath9-gRNA3-Fw | ctcgGGGAATGTACTCCAACATCC | |
| Mpsabath9-gRNA3-Rv | aaacGGATGTTGGAGTACATTCCC | |
| Mpsabath10-gRNA1-Fw | ctcgGTTACGCTGAGAACTCAAAC | Genome editing of Mp*SABATH10* |
| Mpsabath10-gRNA1-Rv | aaacGTTTGAGTTCTCAGCGTAAC | |
| Mpsabath10-gRNA2-Fw | ctcgGAACTTTCTGGATGCTCGGG | |
| Mpsabath10-gRNA2-Rv | aaacCCCGAGCATCCAGAAAGTTC | |
| Mpsabath10-gRNA3-Fw | ctcgTACGAGCAAGCCCTCCTGGA | |
| Mpsabath10-gRNA3-Rv | aaacTCCAGGAGGGCTTGCTCGTA | |
| Mpsabath11-gRNA1-Fw | ctcgAAGTCCAACTTTCCGGTAGA | Genome editing of Mp*SABATH11* |
| Mpsabath11-gRNA1-Rv | aaacTCTACCGGAAAGTTGGACTT | |
| Mpsabath11-gRNA2-Fw | ctcgAGCGTGTTGAAATCGGTGTT | |
| Mpsabath11-gRNA2-Rv | aaacAACACCGATTTCAACACGCT | |
| Mpsabath11-gRNA3-Fw | ctcgAATCTTCCGGATTGCTCGTC | |
| Mpsabath11-gRNA3-Rv | aaacGACGAGCAATCCGGAAGATT | |

| Name | Oligo Sequence (5′ -> 3′) | Used for |
|---|---|---|
| MpCPS-NL1-OligoA | ctcgATCAACCTTACGAACCGGAC | Genome editing of Mp*CPS* |
| MpCPS-NL1-OligoB | aaacGTCCGGTTCGTAAGGTTGAT | |
| MpCPS-NL2-OligoA | ctcgGATAACTGCCACAGCGAAGC | |
| MpCPS-NL2-OligoB | aaacGCTTCGCTGTGGCAGTTATC | |
| MpCPS-NR1-OligoA | ctcgTTCGGGTACAAGGGTTTGGA | |
| MpCPS-NR1-OligoB | aaacTCCAAACCCTTGTACCCGAA | |
| MpCPS-NR2-OligoA | ctcgGAATGTCTAGTACGGAGCTT | |
| MpCPS-NR2-OligoB | aaacAAGCTCCGTACTAGACATTC | |
| MpCPS-gt-F | GGAACCTATCCGGGGATCCT | Genotyping of Mp*cps*$^{ld}$ |
| MpCPS-gt-R | ATGTGACGTTCGTTTGCTGC | |
| Mapoly0140s0010-gRNA1-F | ctcgGATCATGGCTTTTCTCCCGC | Genome editing of Mp*KOL1* |
| Mapoly0140s0010-gRNA1-R | aaacGCGGGAGAAAAGCCATGATC | |
| Mapoly0140s0010-gRNA2-F | ctcgGATGCTCGCTCCATAAAAAC | |
| Mapoly0140s0010-gRNA2-R | aaacGTTTTTATGGAGCGAGCATC | |
| Mapoly0140s0010-gRNA3-F | ctcgATCCGACAAATAATGTTTGT | |
| Mapoly0140s0010-gRNA3-R | aaacACAAACATTATTTGTCGGAT | |
| Mapoly0140s0010-gRNA4-F | ctcgGTTCAGTTTAGAAACCCTCC | |
| Mapoly0140s0010-gRNA4-R | aaacGGAGGGTTTCTAAACTGAAC | |
| Dseq-KOL1-gRNA1~4F | GGATTGATGTACTTGACGAG | Genotyping of Mp*kol1*$^{ld}$ |
| Dseq-KOL1-gRNA1~4R | TTCGGCCTGAAGTCTAAGAG | |
| MpSABATH2_genome_primer1_CACC | caccAAGAGTTCCGCCCAACCGTAAAAG | Cloning of Mp*SABATH2* genomic region |
| MpSABATH2_genome_primer2 | GTCTGCAACAAAAGGACGAG | |
| mCitrine-5end+GGSG-F | GGAGGTAGTGGAATGGTGTCTAAGGGTGAGGA | Insertion of mCitrine to the C-term of Mp*SABATH2* CDS |
| mCitrine-3end-R | CTTGTAAAGCTCATCCATTC | |
| MpSABATH2_C_mCitrine3end-F | GATGAGCTTTACAAGTGAGCCCGCCCGGTCGAATT | |
| MpSABATH2_C_EGFP5end+GGSG | CATTCCACTACCTCCTTTTTTGATCAGGACGGCCA | |
| mCitrine-5end-F | ATGGTGAGCAAGGGCGAGGA | Insertion of mCitrine to the N-term of Mp*SABATH2* CDS |
| mCitrine-3end+GGSG-R | TCCACTACCTCCCTTGTACAGCTCGTCCATGC | |
| MpSABATH2_N_mCitrine-5end-R | GCCCTTGCTCACCATGACTGATACTACAGCTCTCAGAC | |
| MpSABATH2_N_mCitrine-3end+GGSG-F | AAGGGAGGTAGTGGAATGACTGCGGGAGTGCAGAG | |

**Supplemental Table S3**. Sources of genomes used in the phylogenetic analysis

| Clade | Species | Reference | Website |
|---|---|---|---|
| Chlorophytes | *Chlamydomonas reinhardtii* | https://doi.org/10.1093/plcell/koac347 | https://phytozome-next.jgi.doe.gov/info/CreinhardtiiCC_4532_v6_1 |
| Streptophyte algae | *Klebsormidium nitens* | https://doi.org/10.1038/ncomms4978 | http://www.plantmorphogenesis.bio.titech.ac.jp/~algae_genome_project/klebsormidium/kf_download.htm |
| Streptophyte algae | *Chara braunii* | https://doi.org/10.1016/j.cell.2018.06.033 | https://phycocosm.jgi.doe.gov/phycocosm/home |
| Streptophyte algae | *Mesotaenium endlicherianum* | https://doi.org/10.1038/s41477-023-01491-0 | https://mesotaenium.uni-goettingen.de/download.html |
| Streptophyte algae | *Spirogloea muscicola* | https://doi.org/10.1016/j.cell.2019.10.019 | https://figshare.com/articles/dataset/Genomes_of_subaerial_Zygnematophyceae_provide_insights_into_land_plant_evolution/9911876 |
| Streptophyte algae | *Zygnema circumcarinatum* | https://doi.org/10.1101/2023.01.31.526407 | https://phycocosm.jgi.doe.gov/UTEX1560 |
| Mosses | *Ceratodon purpureus* | https://doi.org/10.1126/sciadv.abh2488 | https://phytozome-next.jgi.doe.gov/info/CpurpureusGG1_v1_1 |
| Mosses | *Physcomitrium patens* | https://doi.org/10.1111/tpj.13801 | https://phytozome-next.jgi.doe.gov/info/Ppatens_v3_3 |
| Mosses | *Takakia lepidozioides* | https://doi.org/10.1016/j.cell.2023.07.003 | https://www.takakia.com/download.html |
| Liverworts | *Marchantia polymorpha* | https://doi.org/10.1111/tpj.14602 | https://marchantia.info/data/MpTak_v7.1_standard_genome/ |
| Liverworts | *Marchantia quadrata* | https://doi.org/10.1016/j.celrep.2025.115503 | https://phytozome-next.jgi.doe.gov/info/Mquadratavar_MQ_v1_1 |
| Liverworts | *Lunularia cruciata* | https://doi.org/10.1093/gbe/evad014 | https://genomevolution.org/coge/GenomeInfo.pl?gid=64630 |
| Liverworts | *Riccia fluitans* | https://doi.org/10.1038/s41597-025-04373-6 | https://www.ncbi.nlm.nih.gov/bioproject/PRJNA1158334/ |
| Liverworts | *Conocephalum conicum* | https://doi.org/10.1038/s41586-019-1693-2 https://doi.org/10.1093/gigascience/giz126 | https://db.cngb.org/onekp/species/Conocephalum%20conicum |
| Hornworts | *Anthoceros agrestis* | https://doi.org/10.1038/s41477-024-01883-w | https://hornwortbase.org/ftp/ |
| Hornworts | *Anthoceros fusiformis* | https://doi.org/10.1038/s41477-024-01883-w | https://hornwortbase.org/ftp/ |
| Hornworts | *Leiosporoceros dussii* | https://doi.org/10.1038/s41477-024-01883-w | https://hornwortbase.org/ftp/ |
| Lycophytes | *Selaginella moellendorffii* | https://doi.org/10.1126/science.1203810 | https://phytozome-next.jgi.doe.gov/info/Smoellendorffii_v1_0 |
| Lycophytes | *Isoetes taiwanese* | https://doi.org/10.1038/s41467-021-26644-7 | https://genomevolution.org/coge/GenomeInfo.pl?gid=61511 |
| Ferns | *Adiantum capillus-veneris* | https://doi-org/10.1038/s41477-022-01222-x | https://figshare.com/s/47be9fe90124b22d3c0e |
| Ferns | *Azolla filiculoides* | https://doi.org/10.1038/s41477-018-0188-8 | ftp://ftp.fernbase.org/ |
| Ferns | *Salvinia cucullata* | https://doi.org/10.1038/s41477-018-0188-8 | ftp://ftp.fernbase.org/ |
| Ferns | *Alsophila spinulosa* | https://doi.org/10.1038/s41477-022-01146-6 | https://figshare.com/articles/dataset/A_spinulosa_genome_rar/19075346 |
| Gymnosperms | *Ginkgo biloba* | https://doi.org/10.1038/s41477-021-00933-x | https://figshare.com/articles/dataset/annotation_of_Ginkgo_biloba/14759223 |
| Gymnosperms | *Picea abies* | https://doi.org/10.1038/nature12211 | ftp://plantgenie.org/Data/ConGenIE/ |
| Basal Angiosperms | *Amborella trichopoda* | https://doi.org/10.1126/science.1241089 | https://phytozome-next.jgi.doe.gov/info/Atrichopodavar_SantaCruz_75HAP2_v2_1 |
| Basal Angiosperms | *Nymphaea colorata* | https://doi.org/10.1038/s41586-019-1852-5 | https://phytozome-next.jgi.doe.gov/info/Ncolorata_v1_2 |
| Monocots | *Oryza sativa* | http://dx.doi.org/10.1186/1939-8433-6-4 | https://rapdb.dna.affrc.go.jp/download/irgsp1.html |
| Eudicots | *Arabidopsis thaliana* | https://doi.org/10.1111/tpj.13415 | https://phytozome-next.jgi.doe.gov/info/Athaliana_Araport11 |

### Supplemental References


Bowman, J.L., Kohchi, T., Yamato, K.T., Jenkins, J., Shu, S., Ishizaki, K., et al. (2017) Insights into land plant evolution garnered from the *Marchantia polymorpha* genome. *Cell*. 171: 287–304.

Frank, M.H., and Scanlon, M.J. (2015) Transcriptomic Evidence for the Evolution of Shoot Meristem Function in Sporophyte-Dominant Land Plants through Concerted Selection of Ancestral Gametophytic and Sporophytic Genetic Programs. *Mol Biol Evol*. 32: 355–367.

Higo, A., Niwa, M., Yamato, K.T., Yamada, L., Sawada, H., Sakamoto, T., et al. (2016) Transcriptional Framework of Male Gametogenesis in the Liverwort Marchantia polymorpha L. *Plant Cell Physiol*. 57: 325–338.

Hisanaga, T., Fujimoto, S., Cui, Y., Sato, K., Sano, R., Yamaoka, S., et al. (2021) Deep evolutionary origin of gamete-directed zygote activation by KNOX/BELL transcription factors in green plants. *eLife*. 10: e57090.

Julca, I., Ferrari, C., Flores-Tornero, M., Proost, S., Lindner, A.-C., Hackenberg, D., et al. (2021) Comparative transcriptomic analysis reveals conserved programmes underpinning organogenesis and reproduction in land plants. *Nat Plants*. 7: 1143–1159.

Karaaslan, E.S., Wang, N., Faiß, N., Liang, Y., Montgomery, S.A., Laubinger, S., et al. (2020) Marchantia TCP transcription factor activity correlates with three-dimensional chromatin structure. *Nat Plants*. 6: 1250–1261.

Kawamura, S., Romani, F., Yagura, M., Mochizuki, T., Sakamoto, M., Yamaoka, S., et al. (2022) MarpolBase Expression: a web-based, comprehensive platform for visualization and analysis of transcriptomes in the liverwort *Marchantia polymorpha*. *Plant Cell Physiol*. 63: 1745–1755.

Yasui, Y., Tsukamoto, S., Sugaya, T., Nishihama, R., Wang, Q., Kato, H., et al. (2019) GEMMA CUP-ASSOCIATED MYB1, an Ortholog of Axillary Meristem Regulators, Is Essential in Vegetative Reproduction in *Marchantia polymorpha*. *Curr Biol*. 29: 3987-3995.e5.

Zhao, N., Ferrer, J.-L., Ross, J., Guan, J., Yang, Y., Pichersky, E., et al. (2008) Structural, Biochemical, and Phylogenetic Analyses Suggest That Indole-3-Acetic Acid Methyltransferase Is an Evolutionarily Ancient Member of the SABATH Family. *Plant Physiol*. 146: 323–324.

Zubieta, C., Ross, J.R., Koscheski, P., Yang, Y., Pichersky, E., and Noel, J.P. (2003) Structural Basis for Substrate Recognition in the Salicylic Acid Carboxyl Methyltransferase Family. *Plant Cell*. 15: 1704–1716.
